# Slowly evolving dopaminergic activity modulates the moment-to-moment probability of movement initiation

**DOI:** 10.1101/2020.05.13.094904

**Authors:** Allison E. Hamilos, Giulia Spedicato, Ye Hong, Fangmiao Sun, Yulong Li, John A. Assad

**Affiliations:** Department of Neurobiology, Harvard Medical School, Boston, Massachusetts, 02115, USA; State Key Laboratory of Membrane Biology, Peking University School of Life Sciences, Beijing, 100871, P.R. China; Istituto Italiano di Tecnologia, Genova, Italy

## Abstract

Clues from human movement disorders have long suggested that the neurotransmitter dopamine plays a key role in motor control, but how the endogenous dopaminergic system regulates movement is unknown. Here we show dynamic dopaminergic signaling over seconds-long timescales controls movement timing in mice. Animals were trained to initiate licking after a self-timed interval following a start-timing cue. The movement time was variable from trial-to-trial, as expected from previous studies. Surprisingly, dopaminergic signals ramped-up over seconds between the start-timing cue and the self-timed movement, with variable dynamics that predicted the movement time on single trials. Steeply rising signals preceded early lick-initiation, whereas slowly rising signals preceded later initiation. Higher baseline signals also predicted earlier self-timed movements. Optogenetic activation of dopamine neurons during self-timing did not trigger immediate movements, but rather caused systematic early-shifting of movement initiation, whereas inhibition caused late-shifting, as if modulating the probability of movement. Consistent with this view, the dynamics of the endogenous dopaminergic signals quantitatively predicted the moment-by-moment probability of movement initiation on single trials. These results reveal a causal role for dynamic dopaminergic signaling unfolding over seconds in modulating the decision of when to move.

## INTRODUCTION

What makes us move? Empirically, a few hundred milliseconds before movement, thousands of neurons in the motor system suddenly become active in concert, and this neural activity is relayed via spinal and brainstem neurons to recruit muscle fibers that power movement (***Shenoy et al., 2013***). Yet just before this period of intense neuronal activity, the motor system is largely quiescent. How does the brain suddenly and profoundly rouse motor neurons into the coordinated action needed to trigger movement?

In the case of movements made in reaction to external stimuli, activity evoked first in sensory brain areas is presumably passed along to appropriate motor centers to trigger this coordinated neural activity, thereby leading to movement. But humans and animals can also self-initiate movement without overt, external input (***Deecke, 1996; Hallett, 2007; Lee and Assad, 2003; Romo et al., 1992***). For example, while reading this page, you may decide without prompting to reach for your coffee. In that case, the movement cannot be clearly related to an abrupt, conspicuous sensory cue. What “went off” in your brain that made you reach for your coffee at this *particular* moment, as opposed to a moment earlier or later?

Human movement disorders may provide clues to this mystery. Patients and animal models of Parkinson’s Disease experience difficulty self-initiating movements, exemplified by perseveration (***Hughes et al., 2013***), trouble initiating steps when walking (***Bloxham et al., 1984***), and problems timing movements (***Malapani et al., 1998; Meck, 1986, 2006; Mikhael and Gershman, 2019***). In contrast to these self-generated actions, externally cued reactions are often less severely affected in Parkinson’s, a phenomenon sometimes referred to as “paradoxical kinesia” (***Barthel et al., 2018; Bloxham et al., 1984***). For example, patients’ gait can be normalized by walking aids that prompt steps in reaction to visual cues displayed on the ground (***Barthel et al., 2018***).

Because the underlying neuropathophysiology of Parkinson’s includes the loss of dopaminergic neurons (DANs), the symptomatology of Parkinson’s suggests DAN activity plays an important role in deciding when to self-initiate movement. Indeed, pharmacological manipulations of the neurotransmitter dopamine causally and bidirectionally influence movement timing (***Dews and Morse, 1958; Lustig and Meck, 2005; Meck, 1986; Mikhael and Gershman, 2019; Schuster and Zimmerman, 1961***). This can be demonstrated in the context of *self-timed* movement tasks, in which subjects reproduce a target-timing interval by making a movement following a self-timed delay that is referenced to a start-timing cue (***Malapani et al., 1998***). Species across the animal kingdom, from rodents and birds to primates, can learn these tasks and produce self-timed movements that occur, on average, at about the target time, although the exact timing exhibits considerable variability from trial-to-trial (***Gallistel and Gibbon, 2000; Meck, 2006; Mello et al., 2015; Merchant et al., 2013; Rakitin et al., 1998; Remington et al., 2018; Schuster and Zimmerman, 1961; Sohn et al., 2019; Wang et al., 2018***). In such self-timed movement tasks, decreased dopamine availability/efficacy (*e.g*., Parkinson’s, neuroleptic drugs) generally produces late-shifted movements (***Malapani et al., 1998; Meck, 1986, 2006; Merchant et al., 2013***), whereas high dopamine conditions (*e.g*., amphetamines) produce early-shifting (***Dews and Morse, 1958; Schuster and Zimmerman, 1961***).

Although exogenous dopamine manipulations can influence timing behavior, it remains unknown whether endogenous DAN activity is involved in determining when to move. DANs densely innervate the striatum, where they modulate the activity of spiny projection neurons of the direct and indirect pathways, which are thought to exert a push-pull influence on movement centers (***Albin et al., 1989; DeLong, 1990; Freeze et al., 2013; Grillner and Robertson, 2016***). Moreover, phasic bursts of dopaminergic activity have been observed just prior to movement onset (within ∼500 ms) (***Coddington and Dudman, 2018, 2019; da Silva et al., 2018; Dodson et al., 2016; Howe and Dombeck, 2016; Wang and Tsien, 2011***), and dopaminergic signals have been reported to reflect more general encoding of movement kinematics (***Barter et al., 2015; Engelhard et al., 2019; Parker et al., 2016***). However, optogenetic activation of dopamine neurons—within physiological range—does not elicit immediate movements (***Coddington and Dudman, 2018, 2019***). We hypothesized that rather than overtly triggering movements, the ongoing activity of nigrostriatal DANs could influence movement initiation over longer timescales by controlling or modulating the moment-by-moment decision of *when* to execute a planned movement.

To test this hypothesis, we trained mice to make a movement (lick) after a self-timed interval following a start-timing cue. The mice learned the timed interval, but, as observed in other species, the exact timing of movement was highly variable from trial-to-trial, spanning seconds. We exploited this inherent variability by examining how moment-to-moment nigrostriatal DAN signals differed when animals decided to move relatively early versus late. We found that dopaminergic signals “ramped up” during the timing interval, with variable dynamics that were highly predictive of trial-by-trial movement timing, even seconds before the movement occurred. Optogenetic DAN manipulation during the timing interval produced bidirectional changes in the probability of movement timing, with activation causing a bias toward earlier self-timed movements and suppression causing a bias toward later self-timed movements. These combined observations suggest a novel role for the dopaminergic system in movement initiation, wherein slowly evolving signals modulate the moment-to-moment probability of whether a movement will occur.

## RESULTS

We trained head-fixed mice to make self-timed movements to receive juice rewards (***Figure 1A***). Animals received an audio/visual start-timing cue and then had to decide when to first-lick in the absence of further cues. Animals only received juice if they waited a proscribed interval following the cue before making their first-lick (>3.3 s in most experiments). As expected from previous studies, the distribution of first-lick timing was broadly distributed over several seconds, and exhibited the canonical scalar property of timing, as described by Weber’s Law (***Figure 1B*** and ***Figure 1—figure supplement 1A-B***; (***Gallistel and Gibbon, 2000***)). We note this variability in timing was not imposed on the animal by training it to reproduce a variety of target intervals (*e.g.*, 2 *vs.* 5 s), but is rather a natural consequence of timing behavior, even for a single target interval.

**Figure 1.**
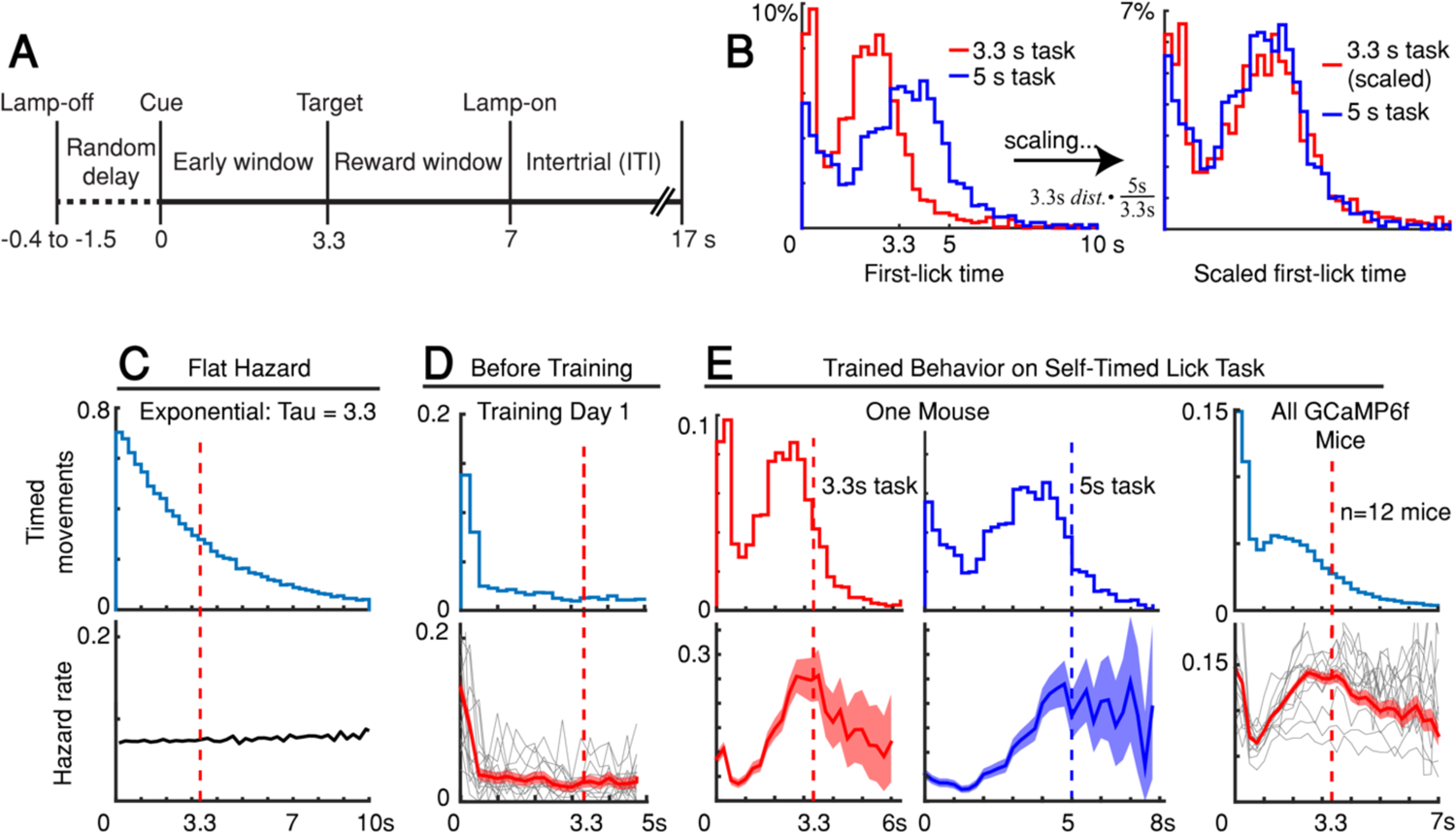
Self-timed movement task. **(A**) Task schematic (3.3 s version shown). **(B)** First-lick timing distributions generated by the same mouse exhibit the scalar property of timing (Weber’s Law). Red: 3.3 s target time (4 sessions); Blue: 5 s target time (4 sessions). For all mice, see ***Figure 1—figure supplement 1B***. (**C-E)** Hazard-function analysis. Time=0 is the start-timing cue; dashed vertical lines are target times. (**C)** Uniform instantaneous probability of movement over time is equivalent to a flat hazard rate (bottom) and produces an exponential first-lick timing distribution (top). (**D)** Before Training: First day of exposure to the self-timed movement task. Top: average first-lick timing distribution across mice; bottom: corresponding hazard functions. Gray traces: single session data. Red traces: average among all sessions, with shading indicating 95% confidence interval produced by 10,000x bootstrap procedure. (**E)** Trained Behavior: Hazard functions (bottom) computed from the first-lick timing distributions for the 3.3 s- and 5 s tasks (top) reveal peaks at the target times. Right: average first-lick timing distribution and hazard functions for all 12 GCaMP6f photometry animals. See also ***Figure 1—figure supplements 1-2*.** Source data: ***Figure 1—source data***.

Our main objective was to exploit the inherent variability in self-timed behavior to examine how differences in neural activity might relate to variability in movement timing. Nonetheless, the trained animals well-understood the timing contingencies of the task. In self-timed movement tasks in which a *single* movement is used to assess timing, the distributions of movement times (in both rodents and monkeys) tend to anticipate the target interval, even at the expense of reward on many trials ***(Eckard and Kyonka, 2018; Kirshenbaum et al., 2008; Lee and Assad, 2003***). In these paradigms, however, once a movement occurs, it removes future opportunities to move, which creates premature “bias” in the raw timing distributions (***Anger, 1956***). To correct this bias, movement times must be normalized by the (ever-diminishing) number of opportunities to move at each timepoint (***Jaldow et al., 1990***). This yields the hazard function (the conditional probability of movement given that movement has not already occurred, as a function of time), which is equivalent to the instantaneous probability of movement. For example, on the first day of training, our animals displayed fairly flat hazard functions, indicating a uniform instantaneous probability of movement over time—*i.e*., the animals did not yet understand the timing contingency (***Figure 1C-D***). However, after training, the hazard function for our animals peaked near the target time (either 3.3 or 5 s), suggesting an accurate latent timing process reflected in the instantaneous movement probability (***Figure 1E***). Mice trained on a variant of the self-timed movement task without lamp-off/on events showed no systematic differences in their timing distributions (***Figure 1—figure supplement 1C***), suggesting that the mice referenced their timing to the start-timing cue rather than the lamp-off event.

When mice were fully trained, we employed fiber photometry to record the activity of genetically-defined DANs expressing the calcium-sensitive fluorophore GCaMP6f (12 mice, substantia nigra pars compacta (SNc); ***Figure 1—figure supplement 2*)**. We controlled for mechanical/optical artifacts by simultaneously recording fluorescence modulation of a co-expressed, calcium-insensitive fluorophore, tdTomato. We also recorded bodily movements with neck-muscle EMG, high-speed video, and a back-mounted accelerometer.

### DAN signals ramp up slowly between the start-timing cue and self-timed movement

DAN GCaMP6f fluorescence typically exhibited brief transients following cue onset and immediately before first-lick onset (***Figure 2A***), as observed in previous studies (***Coddington and Dudman, 2018; da Silva et al., 2018; Dodson et al., 2016; Howe and Dombeck, 2016; Schultz et al., 1997***). However, during the timed interval, we observed slow “ramping up” of fluorescence over seconds, with a minimum after the cue-aligned transient and maximum just before the lick-related transient. The relatively fast intrinsic decay kinetics of GCaMP6f (t_1/2_ <100 ms at 37°; (***Helassa et al., 2016***)) should not produce appreciable signal integration over the seconds-long timescales of the ramps we observed.

**Figure 2.**
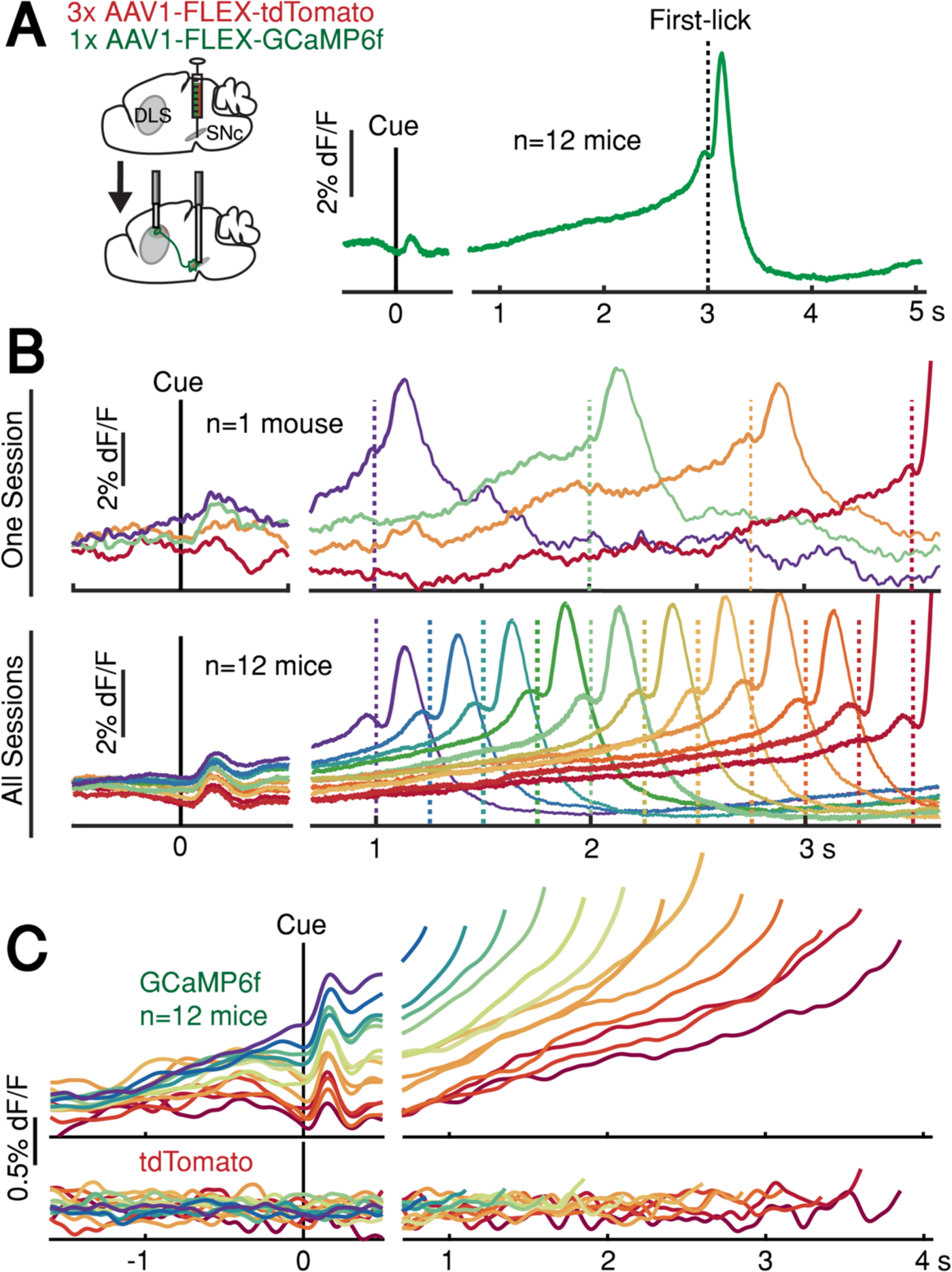
SNc DAN signals preceding self-timed movement. (**A)** Left: surgical strategy for GCaMP6f/tdTomato fiber photometry. Right: average SNc DAN GCaMP6f response for first-licks between 3-3.25 s (12 mice). Data aligned separately to both cue-onset (left) and first-lick (right), with the break in the time axis indicating the change in plot alignment. (**B)** Average SNc DAN GCaMP6f responses for different first-lick times (indicated by dashed vertical lines). **(C)** Comparison of average DAN GCaMP6f and tdTomato responses on expanded vertical scale. Traces plotted up to 150 ms before first-lick. See also ***Figure 2—figure supplements 1-3***. Source data: ***Figure 2—source data***.

We asked whether this ramping differed between trials in which the animal moved relatively early or late. Strikingly, when we averaged signals pooled by movement time, we observed systematic differences in the steepness of ramping that were highly predictive of movement timing (***Figure 2B-C***). Trials with early first-licks exhibited steep ramping, whereas trials with later first-licks started from lower fluorescence levels and rose more slowly toward the time of movement.

The fluorescence ramps terminated at nearly the same amplitude, regardless of the movement time. Ramping dynamics were not evident in control tdTomato signals (***Figure 2C***), indicating that the ramping in the GCaMP6f signals was not an optical artifact. The quantitative relationship between GCaMP6f dynamics and movement time will be addressed in a subsequent section of this paper.

### Higher pre-cue DAN signals are correlated with earlier self-timed movements

In addition to ramping dynamics, average DAN GCaMP6f signals were correlated with first-lick timing even before cue-onset (the “baseline” period, between lamp-off to cue event), with higher baseline fluorescence predicting earlier first-licks (Pearson’s r=-0.89, n=12 mice; ***Figure 2B-C***). Because dF/F correction methods can potentially distort baseline measurements, we rigorously tested and validated three different dF/F methods, and we also repeated analyses with raw fluorescence values compared between pairs of sequential trials with different movement times (***Figure 2—figure supplement 1***; see ***Methods***). All reported results, including the systematic baseline differences, were robust to dF/F correction.

In principle, the amplitude of the baseline signal on a given trial *n* could be related to the outcome of the previous trial or the animal’s behavior during the baseline interval. To test this, we performed 4-way ANOVA on the baseline signal (averaged for each trial between lamp-off and the start-timing cue, n=12 mice), including factors 1) outcome of the previous (*n*-1^th^) trial (rewarded or unrewarded); 2) presence or absence of spontaneous licking during the baseline period; 3) upcoming movement time on trial *n* (categorized as <3.3 s or >3.3 s, to provide a simple binary proxy for movement time); and 4) session number (to account for signal variability across animals and daily sessions). Although previous trial outcome and baseline-licking were significant predictors of baseline amplitude (p<0.01 for both), the upcoming movement time had a significant independent (main) effect (p<10^-5^). This raises the possibility of an additional source of variance in baseline dopaminergic activity that is independent from previous trial events, but potentially influences the upcoming movement time on that trial.

### Movement timing-related ramping dynamics in other dopaminergic areas and striatal dopamine release

We found similar ramping dynamics in SNc DAN axon terminals in the dorsolateral striatum (DLS; ***Figure 2—figure supplement 2A-B*)** at a location involved in goal-directed licking behavior (***Sippy et al., 2015***). Ramping was also present in GCaMP6f-expressing DAN cell bodies in the ventral tegmental area (VTA, ***Figure 2—figure supplement 2C***), reminiscent of mesolimbic ramping signals described in goal-oriented navigation tasks (***Howe et al., 2013; Kim et al., 2019***).

To determine if these movement-timing-related signals are available to downstream targets that may be involved in movement initiation, we monitored dopamine release in the DLS with two complementary florescent dopamine sensors (dLight1.1 and DA_2m_) expressed broadly in striatal cells (***Figure 3* and *Figure 2—figure supplement 2D-E***). The decay kinetics of the two extracellular dopamine sensors differ somewhat (***Patriarchi et al., 2018; Sun et al., 2020***), which we confirmed (dLight1.1 t_1/2_∼75 ms, DA_2m_ t_1/2_∼125 ms; ***Figure 3—figure supplement 1***), yet both revealed similar timing-related ramping dynamics on average (***Figure 3 inset***). These combined data argue that the seconds-long dopaminergic ramping signals were not artifacts of sluggish temporal responses of the various fluorescent sensors and were ultimately expressed as ramp-like increases in dopamine release in the striatum.

**Figure 3.**
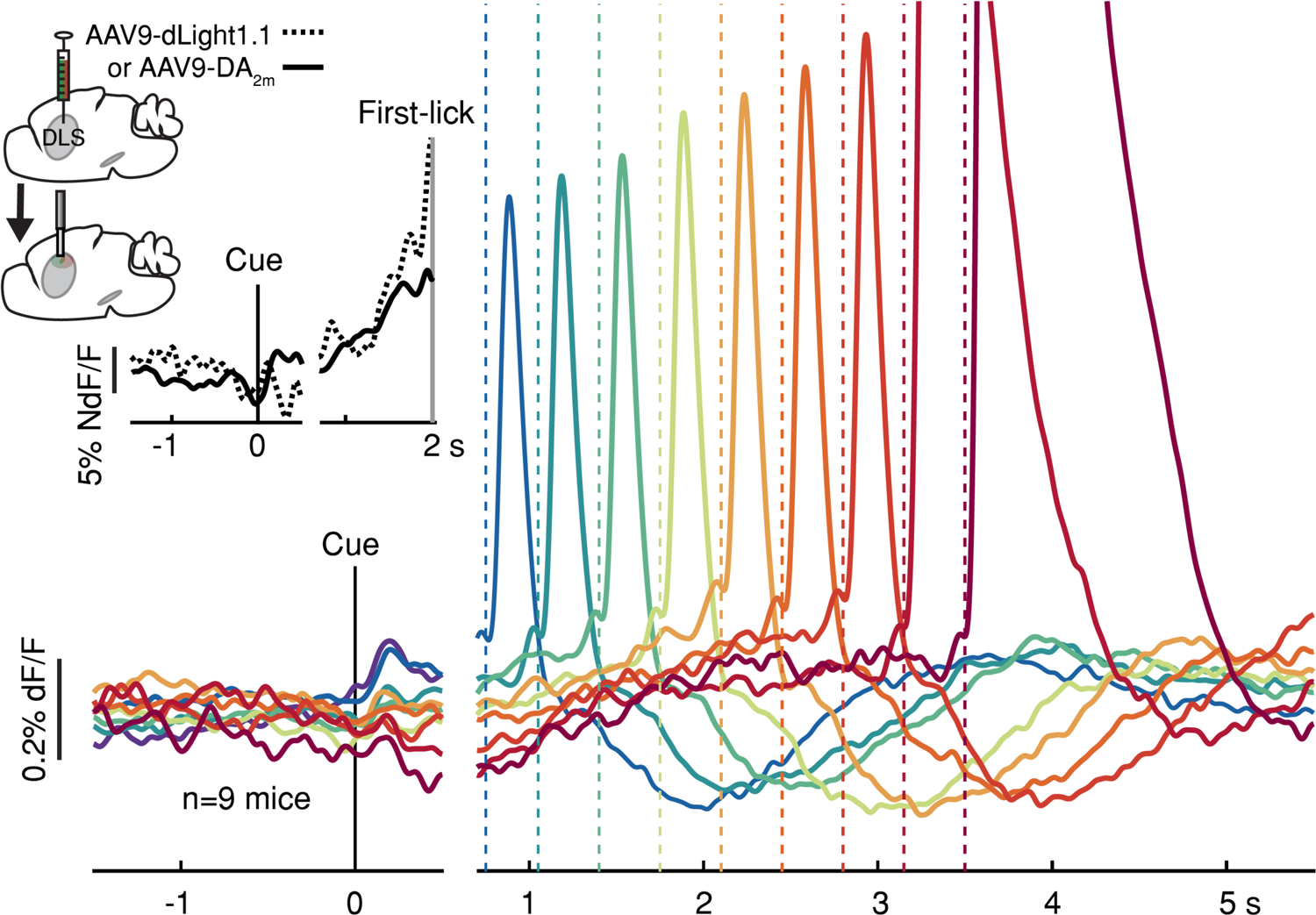
Striatal dopamine release during the self-timed movement task. Photometry signals averaged together from DA_2m_ signals (n=4 mice) and dLight1.1 signals (n=5 mice) recorded in DLS. Axis break and plot alignment as in ***Figure 2***. Dashed lines: first-lick times. Inset, left: surgical strategy. Inset, right: Comparison of dLight1.1 and DA_2m_ dynamics. Expanded vertical scale to show ramping in the average signals for DA_2m_ (solid trace) and dLight1.1 (dashed trace) up until the time of the first-lick (first-lick occurred between 2-3 s after the cue for this subset of the data). See also: ***Figure 3—figure supplement 1***. Source data: ***Figure 3—source data***.

### First-lick timing-predictive DAN signals are not explained by ongoing body movements

The systematic ramping dynamics and baseline differences were not observed in the tdTomato optical control channel nor in any of the other movement-control channels, at least on average (***Figure 4***), making it unlikely that ramping dynamics could have arisen from optical artifacts. Nevertheless, because DANs show transient responses to salient cues and movements (***Coddington and Dudman, 2018; da Silva et al., 2018; Dodson et al., 2016; Howe and Dombeck, 2016; Schultz et al., 1997***), it is possible that fluorescence signals could reflect the superposition of dopaminergic responses to multiple task events, including the cue, lick, ongoing spurious body movements, and hidden cognitive processes like timing. For example, accelerating spurious movements could, in principle, produce motor-related neural activity that “ramps up” during the timed interval, perhaps even at different rates on different trials.

**Figure 4.**
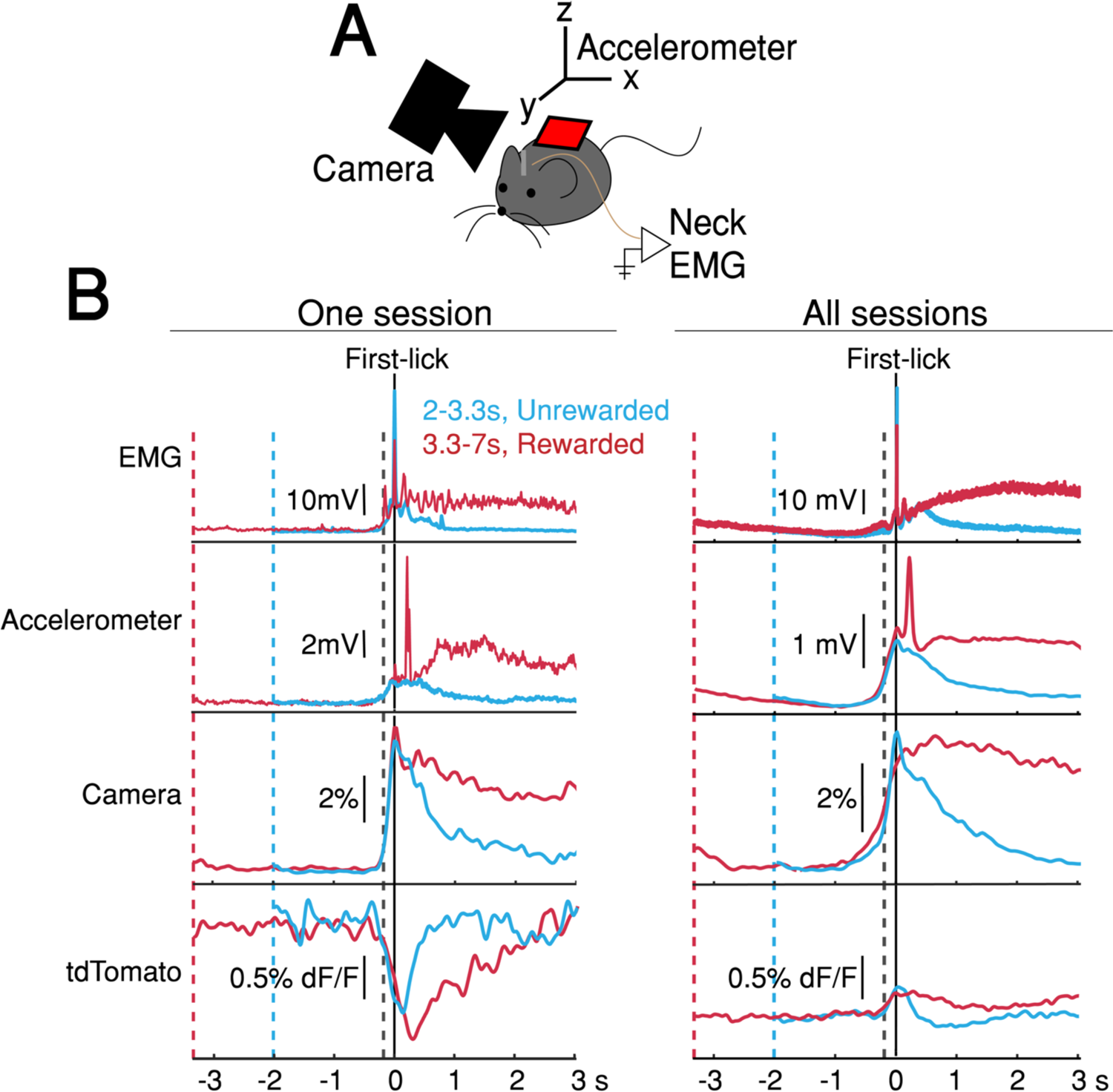
Movement controls reliably detected movements, but there were no systematic differences in movement during the timing interval. **(A)** Schematic of movement-control measurements**. (B)** First-lick-aligned average movement signals on rewarded (red) and unrewarded (blue) trials. Pre-lick traces begin at the nearest cue-time (dashed red, dashed blue). Left: one session; Right: all sessions. Dashed grey line: time of earliest-detected movement on most sessions (150 ms before first-lick). Average first-lick-aligned tdTomato optical artifacts showed inconsistent excursion directions (up/down) even within the same session; signals for each artifact direction shown in ***Figure 4—figure supplement 1***. Source data: ***Figure 4— source data***.

We thus derived a nested generalized linear encoding model of single-trial GCaMP6f signals (***Engelhard et al., 2019; Park et al., 2014; Runyan et al., 2017***), a data-driven, statistical approach designed to isolate and quantify the contributions of task events (timing-independent predictors) from processes predictive of movement timing (timing-dependent predictors; ***Figure 5A-B*** and ***Figure 5—figure supplement 1A-D***). The model robustly detected task-event GCaMP6f kernels locked to cue, lick and EMG/accelerometer events, but these timing-independent predictors alone were insufficient to capture the rich variability of GCaMP6f signals for trials with different first-lick times, especially the timing-dependent ramp-slope and baseline offset (n=12 mice, ***Figure 5C*** and ***Figure 5—figure supplement 1E-G***). In contrast, two timing-dependent predictors robustly improved the model: 1) a baseline offset with amplitude linearly proportional to first-lick time; and 2) a “stretch” feature representing percentages of the timed interval (***Figure 5B-C*** and ***Figure 5—figure supplement 1E***). The baseline offset term fit a baseline level inversely proportional to movement time, and the temporal stretch feature predicted a ramping dynamic from the time of the cue up to the first-lick, whose slope was inversely proportional to first-lick time. Similar results were obtained for SNc DAN axon terminals in the DLS, VTA DAN cell bodies, and extracellular striatal dopamine release (***Figure 5—figure supplement 1H***).

**Figure 5.**
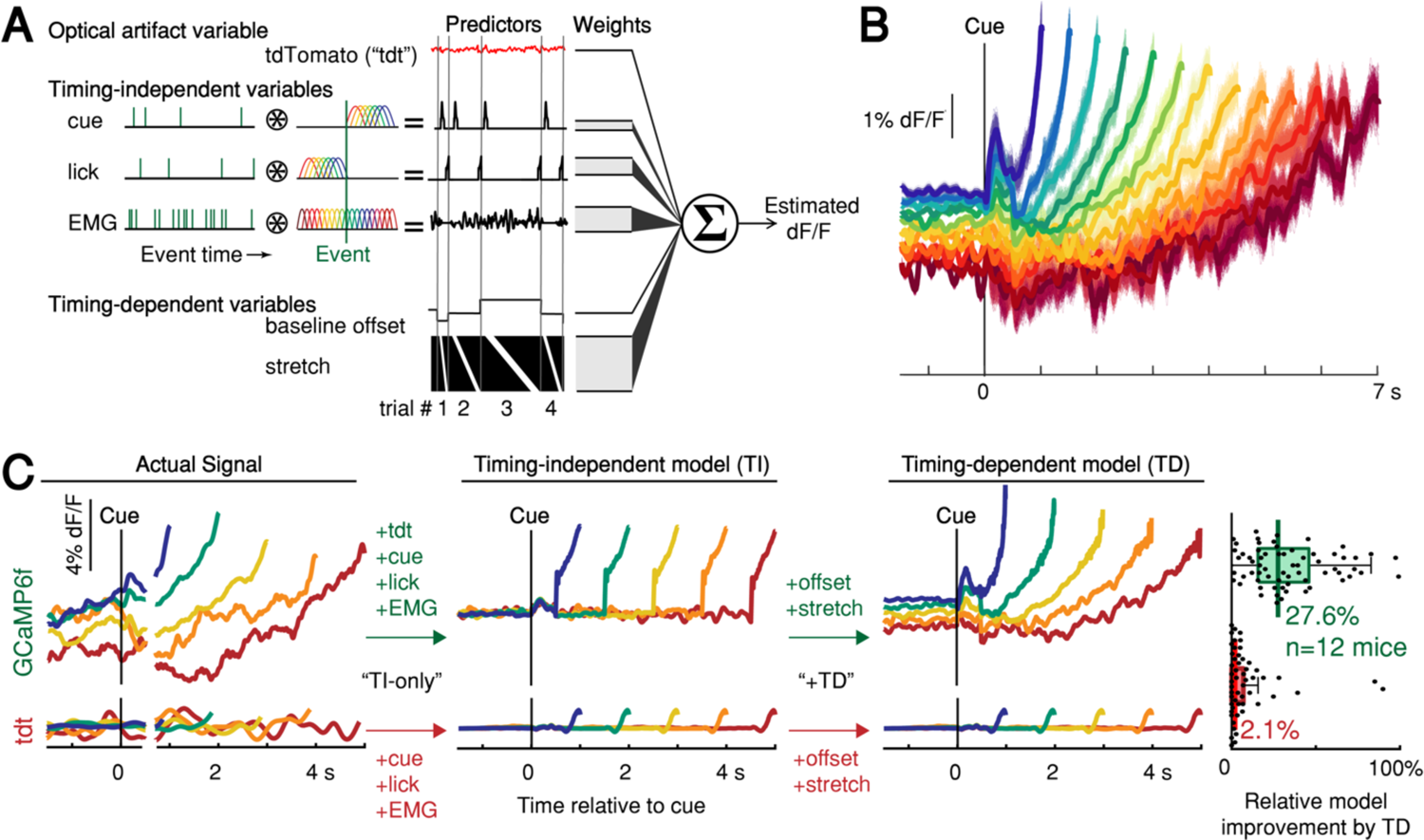
Contribution of optical artifacts, task variables and nuisance bodily movements to SNc GCaMP6f signals. **(A)** Nested encoding model comparing the contribution of timing-independent predictors (TI) to the contribution of timing-dependent predictors (TD). (**B)** Predicted dF/F signal for one session plotted up to time of first-lick. Model error simulated 300x (shading). (**C)** Nested encoding model for one session showing the actual recorded signal (1st panel), the timing-independent model (2nd panel), and the full, timing-dependent model with all predictors (3^rd^ panel). Top: GCaMP6f; Bottom: tdTomato (tdt). Right: relative loss improvement by timing-dependent predictors (grey dots: single sessions, line: median, box: lower/upper quartiles, whiskers: 1.5x IQR). See also ***Figure 5—figure supplement 1***. Source data: ***Figure 5—source data***.

We note that the stretch feature of this GLM makes no assumptions about the underlying shape of the dopaminergic signal; it only encodes percentages of timing intervals to allow for temporal “expansion” or “contraction” to fit whatever shape(s) were present in the data. In particular, the stretch feature cannot produce ramping unless ramping is present in the signal and temporally scales with the length of the interval. Because this feature empirically found a ramp (although not constrained to do so), the stretch aspect indicated that the underlying ramping process took place at different rates for trials with different movement times, at least on average.

In contrast to the GCaMP6f model, when the same GLM was applied to the tdTomato control signal, the timing-independent predictors (which could potentially cause optical/mechanical artifacts—cue onset, first-lick, EMG/accelerometer) improved the model, but timing-dependent predictors did not (***Figure 5C*** and ***Figure 5—figure supplement 1F-H***). In addition, separate principal component (PC) analysis revealed ramp-like and baseline-offset-like components that explained as much as 93% of the variance in DAN signals during the timing interval (mean: 66%, range: 16-93%), but similar PCs were not present when tdTomato control signals were analyzed with PCA (mean variance explained: 4%, range: 1.6-15%, ***Figure 5—figure supplement 2***).

### Single-trial DAN ramping and baseline signals predict movement timing

Given that ramping and baseline-offset signals were not explained by nuisance movements or optical artifacts, we asked whether DAN GCaMP6f fluorescence could predict first-lick timing on single trials. Using a simple threshold-crossing decoding model (***Maimon and Assad, 2006***), we found that single-trial GCaMP6f signals were predictive of first-lick time even for low thresholds intersecting the “base” of the ramp, with the predictive value of the model progressively improving for higher thresholds (n=12 mice; mean R^2^ low: 0.54, mid: 0.71, high: 0.82; analysis for one mouse shown in ***Figure 6A***). We will return to this observation in more detail in the upcoming section on single-trial dynamics (see below).

**Figure 6.**
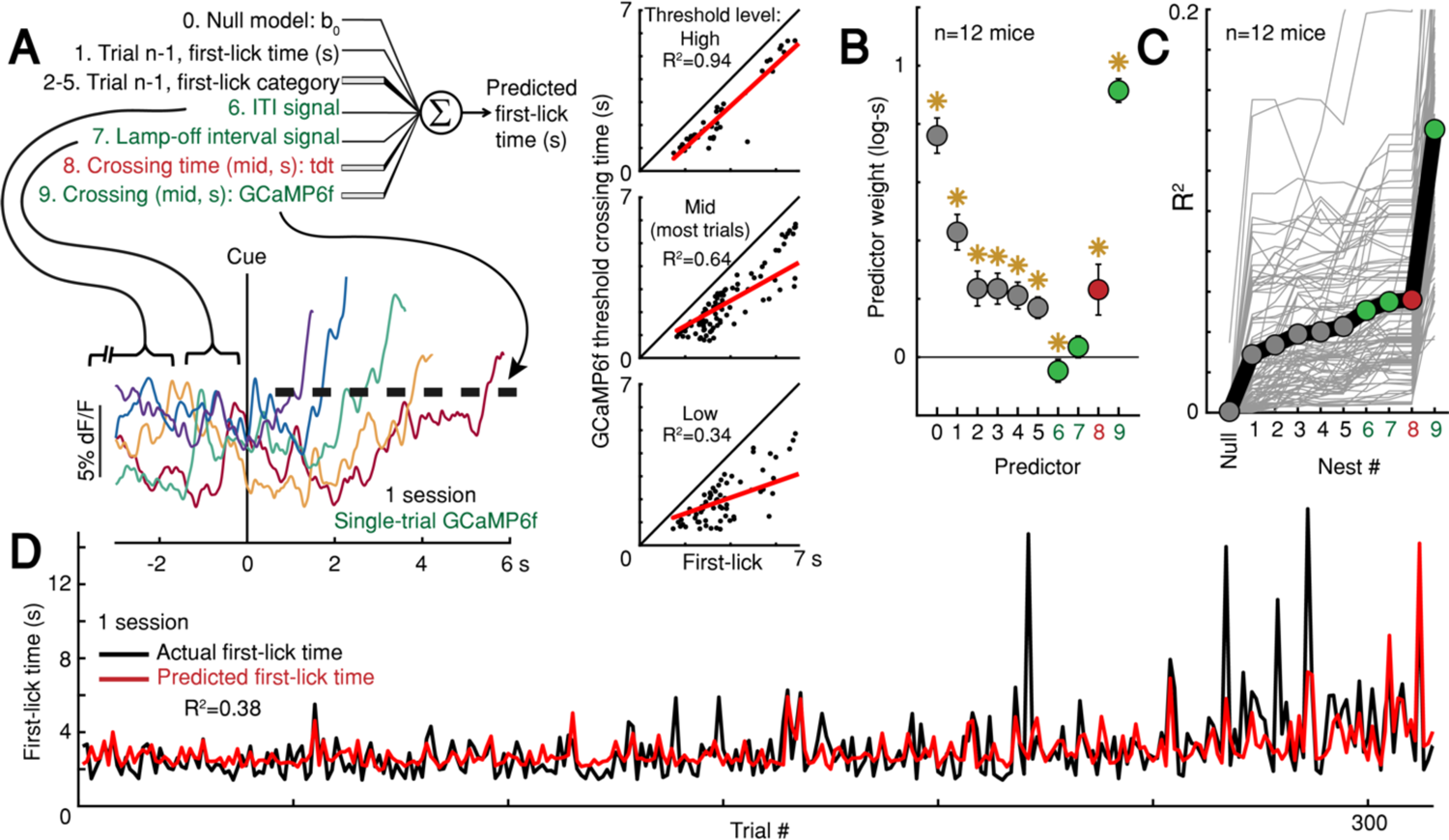
Single-trial DAN signals predict first-lick timing. **(A)** Schematic of nested decoding model. Categories for n-1^th^ trial predictors: 2) reaction, 3) early, 4) reward, 5) ITI first-lick (see ***<u>Methods</u>***). Bottom: single-trial cue-aligned SNc DAN GCaMP6f signals from one session (6 trials shown for clarity). Traces plotted up to first-lick. Right: threshold-crossing model. Low/Mid/High label indicates threshold amplitude. Dots: single trials. **(B)** Model weights. Error bars: 95% CI, *: p<0.05, 2-sided t-test. Numbers indicate nesting-order. **(C)** Variance explained by each model nest. Grey lines: single sessions; thick black line: average. For model selection, see ***Figure 6—figure supplement 1C*. (D)** Predicted *vs*. actual first-lick time, same session as ***6A***. See also ***Figure 6— figure supplements 1-4*.** Source data: ***Figure 6—source data***.

To more thoroughly determine the independent, additional predictive power of DAN baseline and ramping signals over other task variables (*e.g.*, previous trial first-lick time and reward outcome, *etc*.), we derived a nested decoding model for first-lick time (***Figure 6A***). All predictors contributed to the predictive power of the model. However, even when we accounted for the contributions of prior trial history, tdTomato artifacts and baseline GCaMP6f signals, GCaMP6f threshold-crossing time robustly dominated the model and absorbed much of the variance explained by baseline dopaminergic signals, alone explaining 10% of the variance in first-lick time on average (range: 1-27%, ***Figure 6B-D***). Alternate formulations of the decoding model produced similar results (***Figure 6—figure supplement 1***).

### Characterizing single-trial dopaminergic dynamics

Although the threshold-crossing analysis made no assumptions about the underlying dynamics of the GCaMP6f signals on single-trials, in principle, ramping dynamics in *averaged* neural signals could be produced from individual trials with a single, discrete “step” occurring at different times on different trials. Ramping has long been observed in averaged neural signals recorded during perceptual decision tasks in monkeys, and there has been considerable debate over whether single-trial responses in these experiments are better classified as “ramps” or a single “step” (***Latimer et al., 2015, 2016; Shadlen et al., 2016; Zoltowski et al., 2019; Zylberberg and Shadlen, 2016)***. It has even been suggested that different sampling distributions can produce opposite model classifications in ground-truth synthetic signals (***Chandrasekaran et al., 2018***).

We attempted to classify single-trial dynamics as a discrete stepping or ramping process with hierarchical Bayesian models implemented in probabilistic programs (***Figure 6—figure supplement 2A-B***). However, like the perceptual decision-making studies, we also found ambiguous results, with about half of single-trials best classified by a linear ramp and half best classified by a discrete step dynamic (***Figure 6—figure supplement 2C***). Nonetheless, three separate lines of evidence suggest that single trials are better characterized by slowly-evolving ramps:

First, the relationship of threshold-crossing time to first lick time is different for the step *vs.* ramp models when different threshold levels are sampled (***Maimon and Assad, 2006***), as schematized in ***Figure 6—figure supplement 3A***. Increasing slope of this relationship is consistent with ramps on single trials, but not with a discrete step, which would be expected to have the same threshold-crossing time regardless of threshold level (***Figure 6—figure supplement 3B).*** We found that the slope of this relationship increased markedly as the threshold level was increased, consistent with the ramp model (n=12 mice, mean slope low: 0.46, mid: 0.7, high: 0.82, ***Figure 6—figure supplement 3C***).

Second, if single trials involve a step change occurring at different times from trial-to-trial, then aligning trials on that step should produce a clear step on average (rather than a ramp) (***Latimer et al., 2015***). We thus aligned single-trial GCaMP6f signals according to that optimal step position determined from a Bayesian step model fit for each trial and then averaged the step-aligned signals across trials. The averaged signals did not resemble a step function, but rather yielded a sharp transient superimposed on a “background” ramping signal (***Figure 6—figure supplement 4A***). Step-aligned tdTomato and EMG averages showed a small inflection at the time of the step, but neither signal showed background ramping. This suggests that the detected “steps” in the GCaMP6f signals were likely transient movement artifacts superimposed on the slower ramping dynamic rather than *bona fide* steps.

Third, the ideal step model holds that the step occurs at different times from trial-to-trial, producing a ramping signal when trials are averaged together. In this view, the trial-by-trial variance of the signal should be maximal at the time at which 50% of the steps have occurred among all trials, and the signal should be minimal at the beginning and end of the interval (when no steps or all steps have occurred, respectively). We thus derived the optimal step time for each trial using the Bayesian step model, and then calculated variance as a function of time within pools of trials with similar movement times. The signal variance showed a monotonic downward trend during the timed interval, with a minimum variance at the time of movement rather than at the point at which 50% of steps had occurred among trials, inconsistent with the discrete step model (***Figure 6— figure supplement 4B***).

Thus, altogether, we did not find evidence for a discrete step dynamic on single trials; on the contrary, our observations concord with slow ramping dynamics on single trials. Regardless, our GLM movement-time decoding approaches in ***Figure 6*** did not make any assumptions about underlying single-trial dynamics.

### Moment-to-moment DAN activity causally controls movement timing

Because dopaminergic ramping signals robustly predicted first-lick timing and were apparently transmitted via dopamine release to downstream striatal neurons, ramping DAN activity may causally determine movement timing. To test this, we optogenetically activated or inhibited DANs (in separate experiments) on 30% of randomly-interleaved trials (***Figure 7A*** and ***Figure 7—figure supplement 1***). For activation experiments, we chose light levels that elevated DAN activity within the physiological range observed in our self-timed movement task, as assayed by simultaneous photometry in the DLS with a fluorescent sensor of released dopamine (dLight1.1, ***Figure 7— figure supplement 2***). DAN activation significantly early-shifted the distribution of self-timed movements on stimulated trials compared to unstimulated trials (12 mice, p<2.8*10^-26^, 2-sample Kolmogorov-Smirnov (KS) Test), whereas inhibition produced significant late-shifting compared to unstimulated trials (4 mice, p<0.0004, 2-sided KS Test) (***Figure 7B*** and ***Figure 7—figure supplement 3A***). Stimulation of mice expressing no opsin produced no consistent effect on timing (5 mice, p=0.62, 2-sided KS Test). The direction of these effects was consistent across all animals tested in each category (***Figure 7B***). Complementary analysis methods revealed consistent effects (bootstrapped difference in mean first-lick times between categories: p<0.05, ***Figure 7C-D***; bootstrapped comparison of difference in area under the cdf curves: p<0.05, ***Figure 7—figure supplement 3B***; bootstrapped difference in median first-lick times between categories: p<0.05, ***Figure 7—figure supplement 3C***). Similar effects were obtained with activation of SNc DAN axon terminals in the DLS (2 mice, ***Figure 7—figure supplement 3A-B***).

**Figure 7.**
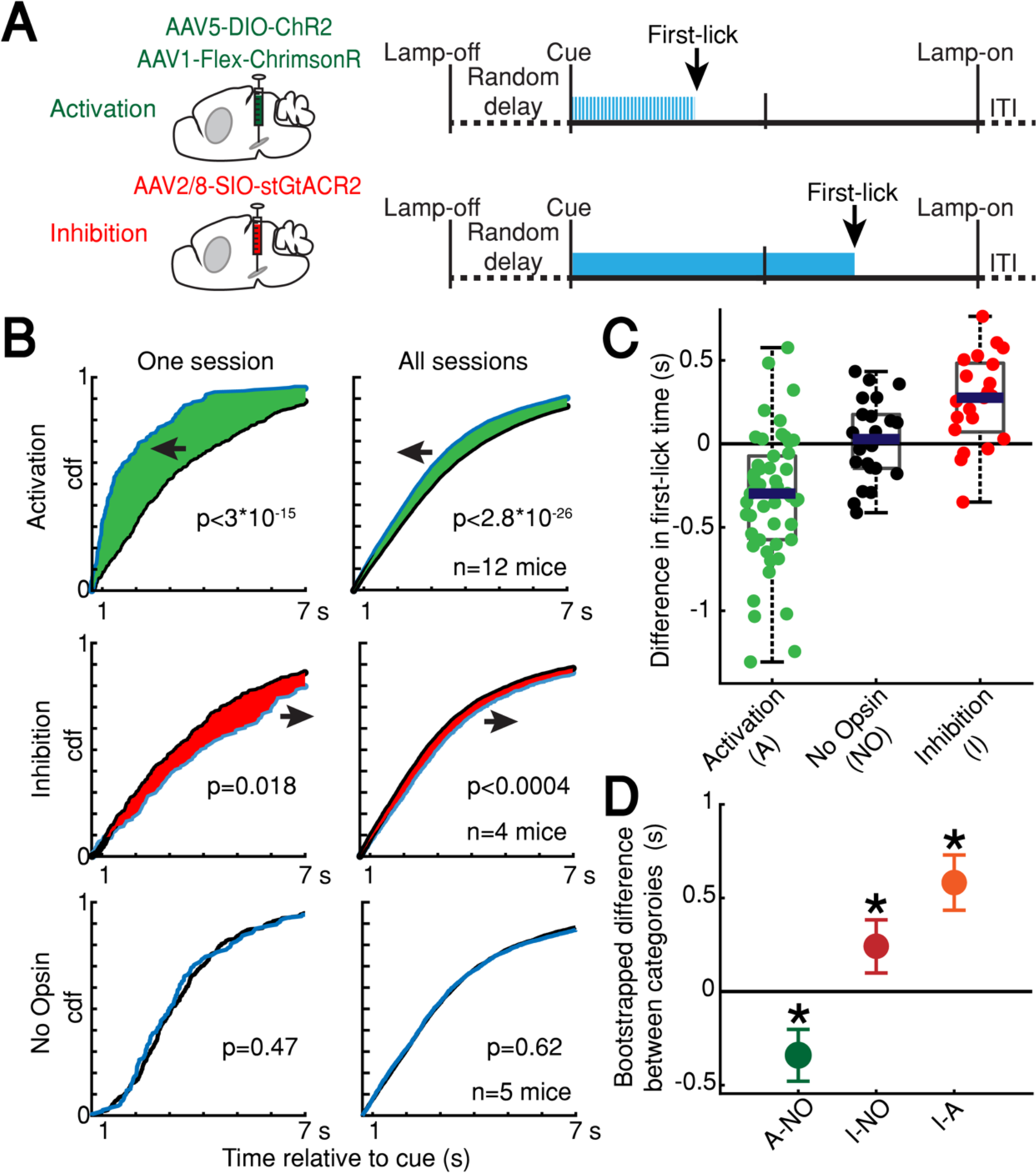
Optogenetic DAN manipulation systematically and bidirectionally shifts the timing of self-timed movements. **(A)** Strategy for optogenetic DAN activation or inhibition. Mice were stimulated from cue-onset until first-lick or 7 s. **(B)** Empirical continuous probability distribution functions (cdf) of first-lick times for stimulated (blue line) versus unstimulated (grey line) trials. Arrow and shading show direction of effect. P-values calculated by Kolmogorov-Smirnov test (for other metrics, see ***Figure 7—figure supplements 1-2***). **(C)** Mean 1,000,000x bootstrapped difference in first-lick time, stimulated-minus-unstimulated trials. Box: upper/lower quartile; line: median; whiskers: 1.5x IQR; dots: single sessions. **(D)** Comparison of mean first-lick time difference across all sessions. Error bars: 95% confidence interval (*: p<0.05, 1,000,000 bootstrapped mean difference in first-lick time between sessions of different stimulation categories). See also ***Figure 7—figure supplements 1-4***. Source data: ***Figure 7—source data***.

Recent studies have shown that physiological ranges of optogenetic DAN activation (as assayed by simultaneous recordings from DANs) fail to elicit overt movements (***Coddington and Dudman, 2018***). We likewise found that optogenetic DAN activation did not elicit immediate licking outside the context of the task (***Figure 7—figure supplement 4A***). Additionally, optogenetic DAN inhibition did not reduce the rate of spontaneous licking outside the context of the task (***Figure 7— figure supplement 4B***). In both cases, we used the same light levels that had elicited the robust shifts in timing behavior during the self-timed movement task. In other control experiments, we purposefully drove neurons into non-physiological activity regimes during the task by applying higher activation light levels. Over-stimulation caused large, immediate, sustained increases in DLS dopamine (***Figure 7—figure supplement 2*)**, comparable in amplitude to the typical reward-related dopamine transients on interleaved, unstimulated trials. These non-physiological manipulations resulted in rapid, nonpurposive body movements and disrupted performance of the task. Together, these results suggest that the optogenetic effects on timing in ***Figure 7*** did not result from direct, immediate triggering or suppression of movement, nor from non-physiological dopamine release due to over-stimulation.

### Linking endogenous DAN signals to the moment-to-moment probability of movement initiation

Optogenetic manipulations of DAN activity in the physiological range appeared to modulate the *probability* of initiating the pre-potent, self-timed movement. Given that endogenous DAN signals increased during the timing interval of the self-timed movement task, we reasoned that the probability of movement should likewise increase over the course of the timed interval. We thus derived a nested probabilistic movement-state decoding model to explore the link between DAN signals and movement propensity (***Figure 8A***). We applied a GLM based on logistic regression, in which we classified each moment of time as either a non-movement (0) or movement (1) state (***Figure 8A-B***), and we examined how well various parameters could predict the probability of transitioning from the non-movement state to the movement state. Unlike the decoding model in **Figure 6**, which considers a single threshold-crossing time, the probabilistic approach takes into account continuous DAN signals. Initial model selection included previous trial history (movement time and reward outcome history) in addition to the DAN GCaMP6f signal, but Bayesian Information Criterion (BIC) analysis indicated that the instantaneous GCaMP6f signal alone was a robustly significant predictor of movement state, whereas previous trial outcomes were insignificant contributors and did not further improve the model (***Figure 8—figure supplement 1***). We thus only considered the DAN GCaMP6f signal as a predictor in subsequent analyses.

**Figure 8.**
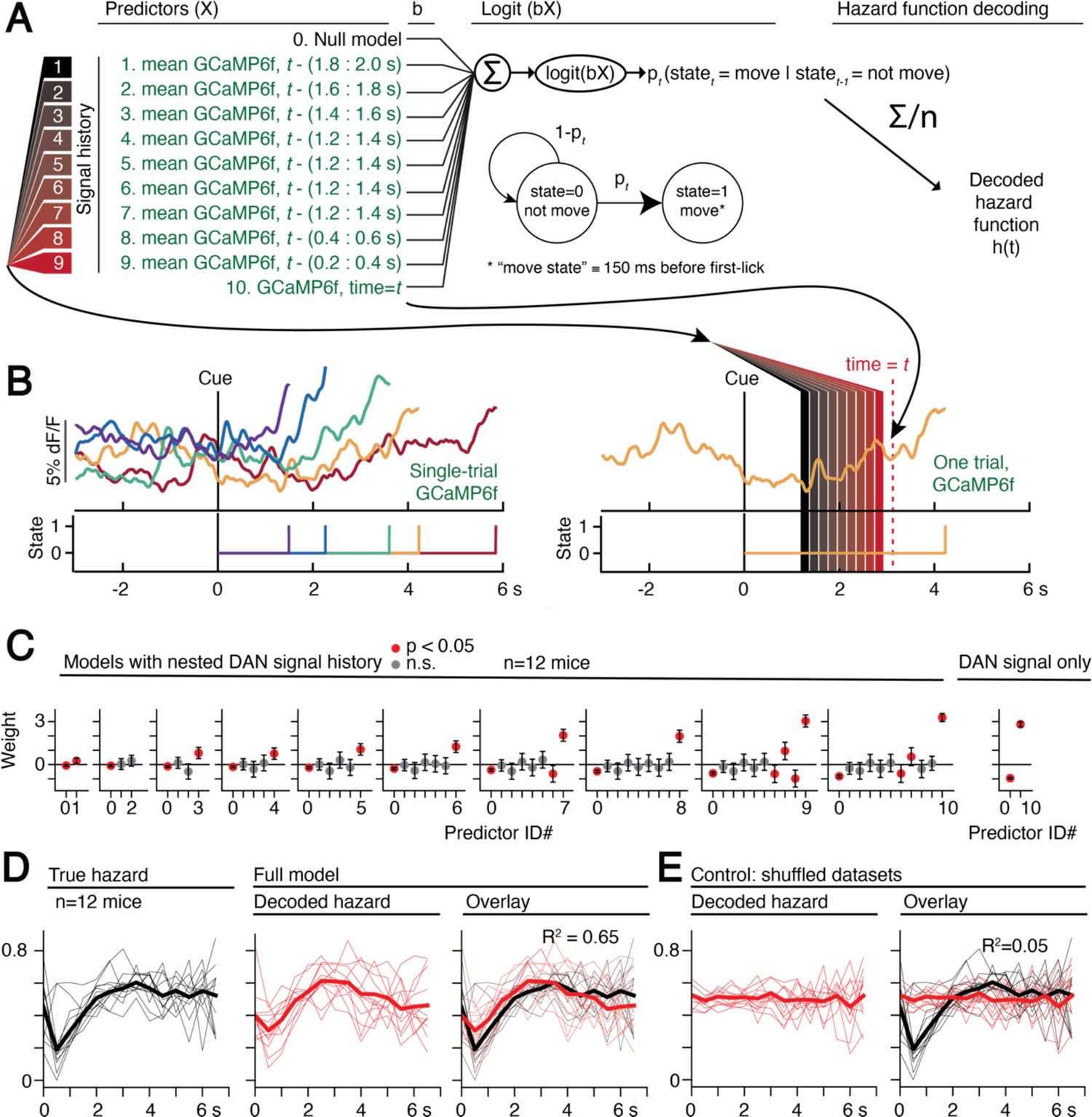
Single-trial dynamic dopaminergic signals predict the moment-to-moment probability of movement initiation. **(A)** Probabilistic movement-state model schematic. **(B)** Single-trial DAN GCaMP6f signals at SNc from one session. First-lick time truncated 150 ms before movement detection to exclude peri-movement signals. Bottom: Movement states for the trials shown as a function of time. Diagram on the right schematizes the model predictors relative to an example time=*t* on a single trial. **(C)** Nested model fitted coefficients. **(D)** Decoded hazard functions from full model (with all 10 predictors). Thick line=mean. n=12 mice. **(E)** Hazard function fitting with shuffled datasets abolished the predictive power of the model (same 12 mice). See also ***Figure 8— figure supplements 1-2***. Source data: ***Figure 8—source data***.

The continuous DAN GCaMP6f signal was indeed predictive of current movement state at any time *t*, and it served as a significant predictor of movement state, up to at least 2 seconds in the past (***Figure 8C***). However, the signals became progressively more predictive of the current movement state as time approached *t*. That is, the dopaminergic signal levels closer to time *t* tended to absorb the behavioral variance explained by more distant, previous signal levels (***Figure 8C***), reminiscent of how threshold crossing time absorbed the variance explained by the baseline dopaminergic signal in the movement-timing decoding model (***Figure 6B-C***). This observation is consistent with a diffusion-like ramping process on single trials, in which the most recent measurement gives the best estimate of whether there will be a transition to the movement state (but is difficult to reconcile with a discrete step process on single trials, consistent with the results in ***Figure 6—figure supplements 3-4***).

We applied the fitted instantaneous probabilities of transitioning to the movement state to derive a fitted hazard function for each behavioral session (***Figure 8D***). The DAN GCaMP6f signals were remarkably predictive of the hazard function, both for individual sessions and on average, explaining 65% of the variance on average (n=12 mice). Conversely, when the model was fit on the same data in which the timepoint identifiers were shuffled, this predictive power was essentially abolished, explaining only 5% of the variance on average (***Figure 8E***).

Together, these results demonstrate that slowly-evolving dopaminergic signals are predictive of the moment-to-moment probability of movement initiation. When combined with the optogenetics results, they argue that dopaminergic signals causally modulate the moment-to-moment probability of the pre-potent movement. In this view, trial-by-trial variability in the DAN signal gives rise to trial-by-trial differences in movement timing in the self-timed movement task.

## DISCUSSION

### A role for slowly ramping DAN activity in the timing of self-initiated movements

We found that both baseline and slowly ramping DAN signals were predictive of the timing of self-initiated movements. A number of studies have reported short-latency (<500 ms) increases in DAN activity following sensory cues and immediately preceding self-initiated movements (***Coddington and Dudman, 2018; da Silva et al., 2018; Dodson et al., 2016; Howe and Dombeck, 2016; Schultz et al., 1997***), similar to the sensory- and motor-related transients we observed within ∼500 ms of the cue and first-lick. However, the ramping DAN signals we observed during self-timing were markedly different. The ramping signal unfolded over *seconds*, preceding the first-lick by as long as 10 s. Furthermore, variations in both ramping dynamics and baseline amplitude predicted the trial-by-trial timing of the first-lick, whether signals were recorded from SNc cell bodies, SNc axon terminals in the DLS, or VTA cell bodies. DAN signals were also reflected in the dynamics of dopamine release in the DLS, indicating availability of this information to downstream striatal effectors.

Optogenetic augmentation and suppression of DAN activity during the timing interval causally altered movement timing. Importantly, optogenetic DAN activation within the physiological range did not evoke immediate movements, consistent with prior work (***Coddington and Dudman, 2018; Lee et al., 2020***). This suggests that, rather than serving as direct drivers of movement, DANs influence *when* movements occur by modulating the moment-by-moment probability of movement. Applying this probabilistic view to the endogenous dopaminergic signals, we found that the ramping dynamics were highly predictive of the moment-by-moment probability of movement (as captured by the hazard function), with DAN signals became progressively better predictors as the time of movement onset approached. These findings suggest that variations in slow DAN dynamics affect movement timing by influencing the moment-to-moment probability of generating a movement.

This view of dopaminergic modulation could be related to classic findings from extrapyramidal movement disorders, in which dysfunction of the nigrostriatal pathway produces aberrations in movement initiation rather than paralysis or paresis (***Bloxham et al., 1984; Fahn, 2011; Hallett and Khoshbin, 1980***). That is, movements do occur in extrapyramidal disorders, but at inappropriate times, either too little/late (*e.g*., Parkinson’s), or too often (*e.g*., dyskinesias). Moreover, based on the deficits observed in Parkinsonian states (*e.g.*, perseveration), this role may extend to behavioral transitions more generally, *e.g*., starting new movements *or* stopping ongoing movements.

### Dopaminergic ramping in other contexts

Previous studies have reported slow ramping dopaminergic signals in certain behavioral contexts, including goal-directed navigation (***Howe et al., 2013***); multi-step tasks culminating in reward (***Hamid et al., 2016; Howard et al., 2017; Mohebi et al., 2019***); and passive observation of dynamic visual cues indicating proximity to reward (***Kim et al., 2019***). It has been proposed that slowly ramping mesolimbic DAN signals could encode increasing value or reward anticipation as animals approach reward (***Hamid et al., 2016; Mohebi et al., 2019***), or alternatively could reflect “ongoing” reward-prediction errors (RPE) (***Kim et al., 2019; Mikhael et al., 2019***). The origin of the ramping signals we observed in the nigrostriatal system are consistent with either value or RPE interpretations. (In a companion theoretical paper (***Hamilos and Assad, 2020***), we examine the possible origins of these dopaminergic signals in value and reward-prediction error computational frameworks (***Kim et al., 2019; Mikhael and Gershman, 2019; Mikhael et al., 2019)***, which offers a reconciliation of apparently contradictory DAN signals reported in the context of a perceptual timing task ***(Soares et al., 2016***)).

Regardless of their origin, it has been unclear how the brain *employs* slowly ramping DAN signals in behavior. Our study moves beyond previous work by finding that trial-by-trial variability in ramping dynamics explains the precise timing of a behavioral output—the self-timed lick—and that optogenetically manipulating SNc DAN activity causally alters the timing of that output. Thus, SNc ramping may not merely encode progress toward a behavioral goal, but could play a causal role in advancing that progress. In this view, ramping DAN signals could be related to, or even drive, ramping signals that have been observed in the motor system in anticipation of self-initiated movements, *e.g.*, readiness potentials in human EEG recordings (Deecke, 1996; Libet et al., 1983).

This view raises the question of whether the slow-timescale DAN signals we observed are unique to the timing requirement of our task or are present before any self-initiated movement. When we averaged DAN signals aligned to “spontaneous” licks during the ITI, we also observed noisy, slow ramping signals building over seconds to the time of the next lick, with a time course related to the duration of the inter-*lick* interval (***Figure 8—figure supplement 2***). This observation raises the possibility that slowly evolving DAN signals may be integral to the generation of self-initiated movements more generally—although our highly trained animals may have also been “rehearsing” timed movements between trials. It would be interesting to see whether slow ramping dynamics predictive of movement timing could be detected in previously published datasets if DAN signals were similarly averaged according to the interval between self-initiated movement bouts.

### Relationship to setpoint and stretching dynamics in other networks during self-timed movement

We found that DAN signals predict movement timing via two low-dimensional signals: a baseline offset and a ramping dynamic that “stretches” depending on trial-by-trial movement timing. Intriguingly, similar stretching of neural responses has been observed before self-timed movement in other brain areas in rats and primates, including the dorsal striatum (***Emmons et al., 2017; Mello et al., 2015; Wang et al., 2018***), lateral interparietal cortex (***Maimon and Assad, 2006***), presupplementary and supplementary motor areas (***Mita et al., 2009***), and dorsomedial frontal cortex (DMFC) (***Remington et al., 2018; Sohn et al., 2019; Wang et al., 2018; Xu et al., 2014***). In the case of DMFC, applying dimensionality reduction to the population responses revealed two lower-dimensional characteristics that resembled our findings in DANs: 1) the speed at which the population dynamics unfolded was scaled (“stretched”) to the length of the produced timing interval (***Wang et al., 2018***), and 2) the population state at the beginning of the self-timed movement interval (“setpoint”) was correlated with the timed interval (***Remington et al., 2018; Sohn et al., 2019***). Recurrent neural network models suggest variation in stretching and setpoint states could be controlled by (unknown) tonic or monotonically-ramping inputs to the cortico-striatal system (***Remington et al., 2018; Sohn et al., 2019; Wang et al., 2018***). We found that DANs exhibit both baseline (*e.g.*, “setpoint”) signals related to timing, as well as monotonically-ramping input during the timing interval. Thus, through their role as diffusely-projecting modulators, DANs could potentially orchestrate variations in cortico-striatal dynamics observed during timing behavior.

### Possible relationship to motivational/movement vigor

In operant tasks in which difficulty is systematically varied over blocks of trials, increased inter-trial dopamine in the nucleus accumbens has been associated with higher average reward rate and with decreased latency to engage in a new trial, suggesting a link between dopamine and “motivational vigor,” the propensity to invest effort in work (***Hamid et al., 2016; Mohebi et al., 2019***). Intriguingly, we observed the *opposite* relationship in the self-timed movement task: periods with higher average reward rates had *lower* average baseline dopaminergic signals and later first-lick times. Moreover, for a given first-lick time (*e.g.*, 3.5-3.75 s), we did not detect differences in baseline (or ramping) signals during periods with different average reward rates, such as near the beginning or end of a session. This difference between the two tasks may be due to their opposing strategic constraints: in the aforementioned experiments, faster trial initiation increased the number of opportunities to obtain reward, whereas earlier first-licks tended to decrease reward acquisition in our self-timed movement task.

The basal ganglia have also been implicated in controlling “movement vigor,” generally referring to the speed, force or frequency of movements (***Bartholomew et al., 2016; Dudman and Krakauer, 2016; Panigrahi et al., 2015; Turner and Desmurget, 2010; Yttri and Dudman, 2016***). The activity of nigrostriatal DANs has been shown to correlate with these parameters during movement bouts and could promote more vigorous movement via push-pull interactions with the direct and indirect pathways (***Barter et al., 2015; da Silva et al., 2018; Mazzoni et al., 2007; Panigrahi et al., 2015***). Movement vigor might also entail earlier self-timed movements, mediated by moment-to-moment increases in dopaminergic activity.

If moving earlier is a signature of greater movement vigor, then earlier self-timed movements might also be executed with greater force/speed. We looked for movement-related vigor signals, examining both the amplitude of lick-related EMG signals and the latency between lick initiation and lick-tube contact. We detected no consistent differences in these force- or speed-related parameters as a function of movement time; on the contrary, the EMG signals were highly stereotyped irrespective of the first-lick time (data not shown). It is possible that vigor might affect movement timing without affecting movement kinematics/dynamics—but, if so, the distinction between “timing” and “vigor” would seem largely semantical.

### Overall view

Lesion and pharmacological studies have long suggested roles for the SNc and dopamine in timing (***Meck, 2006; Merchant et al., 2013***). Broadly speaking, conditions that increase dopamine availability affect timing as if speeding an internal “pacemaker” (***Dews and Morse, 1958; Mikhael and Gershman, 2019; Schuster and Zimmerman, 1961***), whereas conditions that decrease dopamine availability/efficacy generally have the opposite effect (***Malapani et al., 1998; Meck, 1986, 2006; Merchant et al., 2013***). The dopaminergic ramping signals we observed bear some resemblance to Pacemaker-Accumulator models of neural timing, a longstanding conceptual framework for timing behavior in which a hypothetical accumulator signals that an interval has elapsed when it reaches a threshold level (***Gallistel and Gibbon, 2000; Lustig and Meck, 2005; Meck, 2006***).

However, we would suggest a more nuanced view of the role of DANs in self-timed movements. Our optogenetic manipulations indicate that, rather than abruptly triggering movement when threshold is attained, DAN activity modulates the probability of the pre-potent movement. In this view, as DAN activity ramps up, the probability of movement likewise increases, and thus different rates of increase in DAN activity equate to shorter or longer elapsed intervals before movement, on average. This framework leaves open the question of what makes movements “probabilistic.” One possibility is that recurrent cortical-basal ganglia–thalamic circuits could act to generate movements “on their own,” without direct external triggers. By providing crucial modulation of these circuits, DANs could tune the propensity to make self-initiated movements—and pathological loss of DANs could reduce the production of such movements. Future experiments should address how dynamic dopaminergic input influences downstream motor circuits involved in self-initiated movements.

## MATERIALS AND METHODS

### Animals

Adult male and female hemizygous DAT-cre mice **(*Backman et al., 2006***) (B6.SJL-Slc6a3^tm1.1(cre)Bkmm^/J, RRID:IMSR_JAX:020080; The Jackson Laboratory, Bar Harbor, ME) or *wt* C57/b6 mice were used in all experiments (> 2 months old at the time of surgery; median body weight 23.8g, range 17.3-31.9 g). Mice were housed in standard cages in a temperature and humidity-controlled colony facility on a reversed night/day cycle (12 h dark/12 h light), and behavioral sessions occurred during the dark cycle. Animals were housed with enrichment objects provided by the Harvard Center for Comparative Medicine (IACUC-approved plastic toys/shelters, e.g., Bio-Huts, Mouse Tunnels, Nest Sheets, *etc*.) and were housed socially whenever possible (1-5 mice per cage). All experiments and protocols were approved by the Harvard Institutional Animal Care and Use Committee (IACUC protocol #05098, Animal Welfare Assurance Number #A3431-01) and were conducted in accordance with the National Institutes of Health Guide for the Care and Use of Laboratory Animals.

### Surgery

Surgeries were conducted under aseptic conditions and every effort was taken to minimize suffering. Mice were anesthetized with isoflurane (0.5-2% at 0.8 L/min). Analgesia was provided by *s.c*. 5 mg/kg ketoprofen injection during surgery and once daily for 3 d postoperatively (Ketofen, Parsippany, NJ). Virus was injected (50 nL/min) and the pipet remained in place for 10 min before removal. 200 µm, 0.53 NA blunt fiber optic cannulae (Doric Lenses, Quebec, Canada) or tapered fiber optic cannulae (200 µm, 0.60 NA, 2 mm tapered shank, OptogeniX, Lecce, Italy) were positioned at SNc, VTA or DLS and secured to the skull with dental cement (C&B Metabond, Parkell, Edgewood, NY). Neck EMG electrodes were constructed from two Teflon-insulated 32G stainless steel pacemaker wires attached to a custom socket mounted in the dental cement. Sub-occipital neck muscles were exposed by blunt dissection and electrode tips embedded bilaterally.

### Stereotaxic coordinates (from bregma and brain surface)

Viral Injection:

<u>SNc</u>: 3.16 mm posterior, +/- 1.4 mm lateral, 4.2 mm ventral

<u>VTA</u>: 3.1 mm posterior, +/- 0.6 mm lateral, 4.2 mm ventral

<u>DLS</u>: 0 mm anterior, +/- 2.6 mm lateral, 2.5 mm ventral.

Fiber Optic Tips:

<u>SNc/VTA</u>: 4.0 mm ventral (photometry) or 3.9 mm ventral (optogenetics).

<u>DLS</u>: 2.311 mm ventral (blunt fiber) or 4.0 mm ventral (tapered fiber)

### Virus

Photometry:

<u>tdTomato (“tdt”)</u>: AAV1-CAG-FLEX-tdT (UNC Vector Core, Chapel Hill, NC), 100 nL used alone or in mixture with other fluorophores (below), working concentration 5.3*10^12^ gc/mL

<u>gCaMP6f (at SNc or VTA)</u>: 100 nL AAV1.Syn.Flex.GCaMP6f.WPRE.SV40

(2.5*10^13^ gc/mL, Penn Vector Core, Philadelphia, PA). Virus was mixed in a 1:3 ratio with tdt (200 nL total)

<u>DA_2m_ (at DLS)</u>: 200-300 nL AAV9-hSyn-DA4.4(DA2m) (working concentration:

*ca.* 3*\**10^12^ gc/mL, Vigene, Rockville, MD) + 100 nL tdt

<u>dLight1.1 (at DLS)</u>: 300 nL AAV9.hSyn.dLight1.1.wPRE bilaterally at DLS (*ca*.

9.6*10^12^ gc/mL, Children’s Hospital Boston Viral Core, Boston, MA) + 100 nL AAV1.CB7.CI.TurboRFP.WPRE.rBG (*ca.* 1.01*10^12^ gc/mL, Penn Vector Core)

Optogenetic stimulation/inhibition (all bilateral at SNc):

<u>ChR2</u>: 1000 nL AAV5-EF1a-DIO-hChR2(H134R)-EYFP-WPRE-pA (3.2*10^13^ gc/mL,

UNC Vector Core, Chapel Hill, NC)

<u>ChrimsonR +/- dLight1.1</u>: 700 nL AAV1-hSyn-FLEX-ChrimsonR-tdT (4.1*10^12^ gc/mL,

UNC Vector Core, Chapel Hill, NC) +/- 400-550 nL AAV9-hSyn-dLight1.1 bilaterally at DLS (*ca.* 10^13^ gc/mL, Lin Tian Lab, Los Angeles, CA)

<u>stGtACR2</u>: 300 nL 1:10 AAV2/8-hSyn1-SIO-stGtACR2-FusionRed (working concentration 4.7*10^11^ gc/mL, Addgene/Janelia Viral Core, Ashburn, VA)

### Water-deprivation and acclimation

Animals recovered for 1 week postoperatively before water deprivation. Mice received daily water supplementation to maintain ≥80% initial body weight and fed *ad libitum*. Mice were habituated to the experimenter and their health was monitored carefully following guidelines reported previously (***Guo et al., 2014***). Training commenced when mice reached the target weight (∼8-9 d post-surgery).

### Histology

Mice were anesthetized with >400 mg/kg pentobarbital (Somnasol, Henry Schein Inc, Melville, NY) and perfused with 10 mL 0.9% sodium chloride followed by 50mL ice-cold 4% paraformaldehyde in 0.1 M phosphate buffer. Brains were fixed in 4% paraformaldehyde at 4°C for >24 hr before being transferred to 30% sucrose in 0.1 M phosphate buffer for >48 hr. Brains were sliced in 50 µm coronal sections by freezing microtome, and fluorophore expression was assessed by light microscopy. The sites of viral injections and fiber optic placement were mapped with an Allen Mouse Brain Atlas.

### Behavioral rig, data acquisition and analysis

A custom rig provided sensory cues, recorded events and delivered juice rewards under the control of a Teensy 3.2 microprocessor running a custom Arduino state-system behavioral program with MATLAB serial interface. Digital and analog signals were acquired with a CED Power 1400 data acquisition system/Spike2 software (Cambridge Electronic Design Ltd, Cambridge, England). Photometry and behavioral events were acquired at 1,000 Hz; movement channels were acquired at 2,000 Hz. Video was acquired with FlyCap2 or Spinnaker at 30 fps (FLIR Systems, Wilsonville, OR). Data were analyzed with custom MATLAB statistics packages.

### Self-timed movement task

Mice were head-fixed with a juice tube positioned in front of the tongue. The spout was placed as far away from the mouth as possible so that the tongue could still reach it to discourage compulsive licking (***Guo et al., 2014***), ∼1.5 mm ventral and ∼1.5 mm anterior to the mouth. During periods when rewards were not available, a houselamp was illuminated. At trial start, the houselamp turned off, and a random delay ensued (0.4-1.5 s) before a cue (simultaneous LED flash and 3300 Hz tone, 100 ms) indicated start of the timing interval. The timing interval was divided into two windows, early (0-3.333 s in most experiments; 0-4.95 s in others) and reward (3.333-7 s; 4.95-10 s), followed by the intertrial interval (ITI, 7-17 s; 10-20 s). The window in which the mouse first licked determined the trial outcome (early, reward, or no-lick). An early first-lick caused an error tone (440 Hz, 200 ms) and houselamp illumination, and the mouse had to wait until the full timing interval had elapsed before beginning the ITI. Thus there was no advantage to the mouse of licking early. A first-lick during the reward window caused a reward tone (5050 Hz, 200 ms) and juice delivery, and the houselamp remained off until the end of the trial interval. If the timing interval elapsed with no lick, a time-out error tone played (131 Hz, 2 s), the houselamp turned on, and ITI commenced. During the ITI and pre-cue delay (“lamp-off interval”), there was no penalty for licking.

Mice learned the task in 3 stages (***Figure 1—figure supplement 1A***). On the first 1-4 days of training, mice learned a beginner-level task, which was modified in two ways: 1) to encourage participation, if mice did not lick before 5 s post-cue, they received a juice reward at 5 s; and 2) mice were not penalized for licking in reaction to the cue (within 500 ms). When the mouse began self-triggering ≥50% of rewards (day 2-6 of training), the mouse advanced to the intermediate-level task, in which the training reward at 5 s was omitted, and the mouse had to self-trigger all rewards. After completing >250 trials/day on the intermediate task (usually day 4-7 of training), mice advanced to the mature task, with no reaction licks permitted. All animals learned the mature task and worked for ∼400-1,500 trials/session.

### Hazard function correction of survival bias in the timing distribution

The raw frequency of a particular response time in the self-timed movement task is “distorted” by how often the animal has the chance to respond at that time (***Anger, 1956***). This bias was corrected by calculating the hazard function, which takes into account the number of response opportunities the animal had at each timepoint. The hazard function is defined as the conditional probability of moving at a time, *t*, given that the movement has not yet occurred (referred to as “IRT/Op” analysis in the old Differential Reinforcement of Low Rates (DRL) literature). The hazard function was computed by dividing the number of first-movements in each 250 ms bin of the first-lick timing histogram by the total number of first-movements occurring at that bin-time or later—the total remaining “opportunities.”

### Online movement monitoring

Movements were recorded simultaneously during behavior with four movement-control measurements: neck EMG (band-pass filtered 50-2,000 Hz, 60 Hz notch, amplified 100-1,000x), back-mounted accelerometer (SparkFun Electronics, Boulder, CO), high-speed camera (30 Hz, FLIR Systems, Wilsonville, OR), and tdTomato photometry. All control signals contained similar information, and thus only a subset of controls was used in some sessions.

### Photometry

Fiber optics were illuminated with 475 nm blue LED light (Plexon, Dallas, TX) (SNc/VTA: 50 μW, DLS: 35 μW) measured at patch cable tip with a light-power meter (Thorlabs, Newton, NJ). Green fluorescence was collected via a custom dichroic mirror (Doric Lenses, Quebec, Canada) and detected with a Newport 1401 Photodiode (Newport Corporation, Irvine, CA). Fluorescence was allowed to recover ≥1 d between recording sessions. To avoid crosstalk in animals with red control fluorophore expression, the red channel was recorded at one of the 3 sites (SNc, VTA, or DLS, 550 nm lime LED, Plexon, Dallas, TX) while GCaMP6f, dLight1.1 or DA_2m_ was recorded simultaneously only at the other implanted sites.

### dF/F

Raw fluorescence for each session was pre-processed by removing rare singularities (single points >15 STD from the mean) by interpolation to obtain F(t). To correct photometry signals for bleaching, dF/F was calculated as:

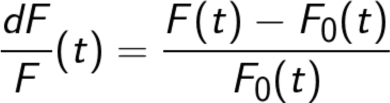

where F_0_(t) is the 200 s moving average of F(t) (***Figure 2—figure supplement 1A***). We tested several other complementary methods for calculating dF/F and all reported results were robust to dF/F method (see ***Methods: dF/F method characterization and validation***). To ensure dF/F signal processing did not introduce artifactual scaling or baseline shifts, we also tested several complementary techniques to isolate undistorted F(t) signals where possible and quantified the amount of signal distortion when perfect isolation was not possible (see ***Methods: dF/F method characterization and validation***, below, and ***Figure 2—figure supplement 1C***).

### dF/F method characterization and validation

dF/F calculations are intended to reduce the contribution of slow fluorescence bleaching to fiber photometry signals, and many such methods have been described (***Kim et al., 2019; Mohebi et al., 2019; Soares et al., 2016***). However, dF/F methods have the potential to introduce artifactual distortion when the wrong method is applied in the wrong setting. Thus, to derive an appropriate dF/F method for use in the context of the self-timed movement task, we characterized and quantified artifacts produced by 4 candidate dF/F techniques.

*Detailed description of complementary dF/F methods*.

1. <u>Normalized baseline</u>: a commonly used dF/F technique in which each trial’s fluorescence is normalized to the mean fluorescence during the 5 s preceding the trial.
2. <u>Low-pass digital filter</u>: F_0_ is the low-pass, digital infinite impulse response (IIR)-filtered raw fluorescence for the whole session (implemented in MATLAB with the built-in function *lowpass* with f_c_=5·10^-5^ Hz, steepness=0.95).
3. <u>Multiple baseline</u>: a variation of Method 1, in which each trial’s fluorescence is normalized by the mean fluorescence during the 5 s preceding the current trial, as well as 5 trials before the current trial and 5 trials after the current trial.
4. <u>Moving average</u>: F_0_ is the 200 s moving average of the raw fluorescence at each point (100 s on either side of the measured timepoint).

Although *normalized baseline* (Method 1) is commonly used to correct raw fluorescence signals (F) for bleaching, this technique assumes that baseline activity has no bearing on the trial outcome; however, because the mouse decides when to move in the self-timed movement task, it is possible that baseline activity may differ systematically with the mouse’s choice on a given trial. Thus, normalizing F to the baseline period would obscure potentially physiologically-relevant signals. More insidiously, if baseline activity *does* vary systematically with the mouse’s timing, normalization can also introduce substantial amplitude scaling and y-axis shifting artifacts when correcting F with this method (***Figure 2—figure supplement 1C***, middle panels). Thus, Methods 2-4 were designed and optimized to isolate photometry signals minimally distorted by bleaching signals and systematic baseline differences during the self-timed movement task. Methods 2-4 produced the same results in all statistical analyses, and the moving average method is shown in all figures.

## Isolating minimally-distorted photometry signals with paired trial analyses of raw fluorescence

Although slow bleaching prevents comparison of raw photometry signals (F) at one time in a behavioral session with those at another time, the time-course of appreciable bleaching was slow enough in the reported behavioral sessions that minimal bleaching occurred over the course of 3 trials (∼1 min, ***Figure 2—figure supplement 1A***). Thus, to observe the most minimally-distorted photometry signals possible, we compared F between *pairs of consecutive* trials (***Figure 2—figure supplement 1B-C***). We compared F baseline signals between all paired trials in which an early trial (unrewarded first-lick between 0.7-2.9 s; abbreviated as “E”) was followed by a rewarded trial (first-lick between 3.4-7 s; abbreviated as “R”); this two-trial sequence is thus referred to as an “ER” comparison. To ensure systematic differences did not result from subtle bleaching in the paired-trial interval, we reversed the ordering contingency and also compared all Rewarded trials preceding Early trials (“RE” comparison). The same systematic relationship between baseline signals and first-lick time was found for paired trials analyzed by raw F (***Figure 2—figure supplement 1C, left panels***).

## Quantification of artifactual amplitude scaling/baseline shifts introduced by dF/F processing

Each Candidate dF/F Method was applied to the same Paired Trial datasets described above. The resulting paired-fluorescence datasets were normalized after processing (minimum dF/F=0, maximum=1). The amount of distortion introduced by dF/F was quantified with a Distortion Index (DI), which was calculated as:

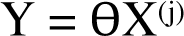

where F(t) and dF/F(t) are the normalized, paired-trial raw fluorescence signal or dF/F signal at time *t*, respectively. *t* spanned from the beginning of the n-1^th^ trial (−20 s) to the end of the n^th^ trial (20 s), aligned to the cue of the n^th^ trial (***Figure 2—figure supplement 1C, bottom panels***). The DI shown in plots has been smoothed with a 200 ms moving average kernel for clarity.

As expected, normalizing fluorescence to the baseline period (*normalized baseline*) erased the correlation of baseline dF/F signals with first-lick time (***Figure 2—figure supplement 1C***, middle panels). More insidiously, this also resulted in distortion of GCaMP6f dynamics *during* the timing interval, evident in the diminished difference between E-signals compared to R-signals relative to the shapes observed in the raw fluorescence paired-trial comparison (***Figure 2—figure supplement 1C, middle-bottom panel***). However, dF/F Methods 2-4 visually and quantitatively recapitulated the dynamics observed in the raw fluorescence comparison (***Figure 2—figure supplement 1C, right panels***).

These results were corroborated by time-in-session permutation tests in which datasets for single sessions were divided into thirds (beginning of session, middle of session, and end of session). The differences between baseline and ramping dynamics observed in whole-session averages were present even within these shorter blocks of time within the session (*i.e*., faster ramping and elevated baseline signals on trials with earlier self-timed licks). Furthermore, permutation tests in which the block identity (begin, middle, end) was shuffled showed that this pattern held when trials with earlier first-licks from the end of the session were compared with trials with later first-licks from the beginning of the session (and *vice versa*).

## Normalized dF/F for comparing dopamine sensor signals

DA_2m_ was about twice as bright as dLight1.1, and thus generally yielded larger and less noisy dF/F signals. To compare the two extracellular dopamine sensors in the same plot, dF/F was normalized for each signal by the amplitude of its lick-related transient. dF/F was calculated as usual, and then the mean baseline-to-transient peak amplitude was measured for trials with first-licks occurring between 2-3 s. Percentage NdF/F is reported as the percentage of this amplitude.

## Dopamine sensor kinetics

dLight1.1 is an extracellular dopamine sensor derived from the dopamine-1-receptor, and has fast reported kinetics: rise t_1/2_ = 9.5 ± 1.1 ms, decay t_1/2_ = 90 ± 11 ms (***Patriarchi et al., 2018***). DA_2m_ is a new extracellular dopamine indicator derived from the dopamine-2-receptor, which provides brighter signals. DA_2m_ signals have been reported to decay slowly in slice preparations but are much faster *in vivo*, presumably because endogenous dopamine-clearance mechanisms are preserved: reported rise t_1/2_ ∼50 ms, decay t_1/2_ ∼360 ms in freely behaving mice; decay t_1/2_ ∼190 ms in head-fixed *drosophila* (***Sun et al., 2020***). To estimate the dopamine-sensor kinetics in our head-fixed mice, we examined the phasic fluorescence transient occurring on unrewarded first-licks (0.5-3.3 s), which showed a stereotyped fast rise and decay with both sensors (***Figure 2—figure supplement 2D-E***). While the transient was somewhat complex (reminiscent of phasic burst-pause responses sometimes observed for movement-related DAN activity (***Coddington and Dudman, 2018, 2019***), we measured the time for average fluorescence to decay from the peak of the transient to half the baseline-to-peak amplitude. We found decay t_1/2_∼75 ms for dLight1.1 and t_1/2_∼125 ms for DA_2m_ (***Figure 3—figure supplement 1***). Given that the dopaminergic ramping signals in our study evolved over several seconds, the kinetics of both dopamine sensors are thus fast enough that they should not have caused appreciable distortion of the slow ramping dynamics.

## Pearson’s correlation of baseline signals to first-lick time

The mean SNc GCaMP6f signal during the minimum lamp-off interval (−0.4 s to 0 s, the cue-time) was compared to the first-lick time for pooled trials in ***Figure 2C*** by calculating the Pearson correlation coefficient. There were at least 700 trials in each pooled set of trials (0.75-4 s included).

## DAN signal encoding model

To test the independent contribution of each task-related input to the photometry signal and select the best model, we employed a nested fitting approach, in which each dataset was fit multiple times (in “nests”), with models becoming progressively more complex in subsequent nests. The nests fit to the GCaMP6f photometry data employed the inputs X^(j)^ at each *j*^th^ nest:

Null Model: X^(0)^ = x_0_
Nest 1: X^(1)^ = X^(0)^ + tdTomato (tdt)
Nest 2: X^(2)^ = X^(1)^ + cue + first-lick
Nest 3: X^(3)^ = X^(2)^ + EMG/accelerometer
Nest 4: X^(4)^ = X^(3)^ + time-dependent baseline offset
Nest 5: X^(5)^ = X^(4)^ + stretch representing percentages of interval

Overfitting was penalized by ridge regression, and the optimal regularization parameter for each nest was obtained by 5-fold cross-validation to derive the final model fit for each session. Model improvement by each input was assessed by the percentage loss improvement at the nest where the input first appeared compared to the prior nest. The loss improvement of Nest 1 was compared to the Null Model (the average of the photometry timeseries). The nested model of tdt control photometry signals was the same, except Nest 1 was omitted.

The GLM for each nest takes the form:

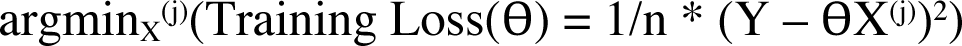

Where Y is the *1*x*n* vector of the photometry signal across an entire behavioral session (*n* is the total number of sampled timepoints); X^(j)^ is the *d*x*n* design matrix for nest *j*, where the rows correspond to the *d_j_* predictors for nest *j* and the columns correspond to each of the *n* sampled timepoints of Y; and ϴ is the *d*x*1* vector of fit weights.

Y is the concatenated photometry timeseries taken from trial start (lights off) to the time of first lick. Because of day-to-day/mouse-to-mouse variation (ascribable to many possible sources, *e.g.*, different neural subpopulations, expression levels, behavioral states, *etc*.), each session was fit separately.

The *d_j_* design matrix predictors were each scaled (maximum amplitude 1) and grouped by input to the model. The timing-independent inputs were: 1. Null offset (x_0_, 1 predictor), 2. tdt (1 predictor), 3. cue (24 predictors), 4. first-lick (28 predictors), and 5. EMG/accelerometer (44 predictors). The timing-dependent inputs were: 6. timing-dependent baseline offset (1 predictor), 7. stretch (500 predictors).

To reduce the number of predictors, the cue, first-lick and EMG/accelerometer predictors (***Figure 5—figure supplement 1C***) were composed from sets of basis kernels as described previously (***Park et al., 2014; Runyan et al., 2017***). The cue basis kernels were spaced 0-500 ms post-cue and first-lick basis kernels were spaced −500 ms-0 ms relative to first-lick, the typically-observed windows of stereotypical sensory and motor-related neural responses. For nuisance movements (EMG/accelerometer), events were first discretized by thresholding (***Figure 5— figure supplement 1B***) and then convolved with basis kernels spanning −500 to 500 ms around the event. This window was consistent with the mean movement-aligned optical artifact observed in the tdt channel. The timing-dependent baseline offset was encoded as a constant offset spanning from lamp-off until first-lick, with amplitude taken as linearly proportional to the timed interval on the current trial. The timing-dependent stretch input was composed of 500 predictors, with each predictor containing 1’s tiling 0.05% of the cue-to-lick interval, and 0’s otherwise (***Figure 5— figure supplement 1D***). Importantly, the stretch was not constrained in any way to form ramps.

Basis sets were optimized to minimize Training Loss, as calculated by mean squared error of the unregularized model:

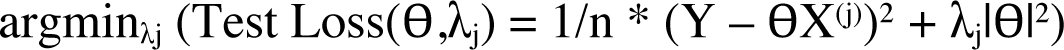

Superfluous basis set elements that did not improve Training Loss compared to the Null Model were not included in the final model. Goodness of the training fit was assessed by Akaike Information Criterion (AIC), Bayesian Information Criterion (BIC), R^2^, and Training Loss. The optimal, regularized model for each nest/session was selected by 5-fold cross-validation in which the regularization parameter, λ_j_, was optimized for minimal average Test Loss:

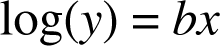

Test Loss for each optimal model was compared across nests to select the best model for each session. Models were refit with the optimal λ_j_ to obtain the final fit.

Model error was simulated 1,000 times by redrawing ϴ coefficients consistent with the data following the method described by Gelman and Hill (***Gelman and Hill, 2006***), and standard errors were propagated across sessions. The absolute value of each predictor was summed and divided by the total number of predictors for that input to show the contribution of the input to the model (***Figure 5—figure supplement 1G***). To simulate the modeled session’s photometry signal for each nest *j*, Yfit was calculated as ϴX^(j)^ and binned by the time of first-lick relative to the cue. The error in the simulation was shown by calculating Yfit_sim_ = ϴ_sim_X^(j)^ for 300 simulated sets of ϴ_sim_.

## Principal component analysis (PCA)

Unsmoothed ramping intervals for photometry timeseries were fit with PCA and reconstructed with the first three principal components (PCs). To derive a PCA fit matrix with ramping intervals of the same number of samples, the length of each trial was scaled up by interpolation to the maximum ramping interval duration:

7 s – 0.7 s cue buffer – 0.6 s first-lick buffer = 5.7 s: 5,700 sample ramping interval Following PC-fitting, datasets were down-sampled to produce a fit of the correct time duration. Trials where the ramping interval was <0.1 s were excluded to exclude noise from down-sampling.

## First-lick time decoding model

A nested, generalized linear model was derived to predict the first-lick time on each trial in a session and quantify the contribution of previous reward history and photometry signals to the prediction. The model was of the form:

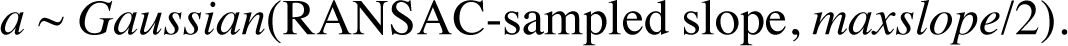

where *y* is the first-lick time, *b* is a vector of fit coefficients and *x* is a vector of predictors. The nested model was constructed such that predictors occurring further back in time (such as reward history) and confounding variables (such as tdt photometry signals) were added first to determine the additional variance explained by predictors occurring closer to the time of first-lick, which might otherwise obscure the impact of these other variables. The predictors, in order of nesting, were:

Nest 0: b0 (Null model, average log-first-lick time)
Nest 1: b1 = b0 + first-lick time on previous trial (trial “n-1”)
Nest 2-5: b2 = b1 + previous trial outcome (1,0)*
Nest 6: b3 = b2 + median photometry signal in 10s window before lamp-off (“ITI”)
Nest 7: b4 = b3 + median photometry signal from lamp-off to cue (“lamp-off interval”)
Nest 8: b5 = b4 + tdt threshold crossing time**
Nest 9: b6 = b5 + GCaMP6f threshold crossing time**

where all predictors were normalized to be in the interval (0,1).

* Outcomes included (in order of nest): Reaction (first-lick before 0.5 s), Early (0.5-3.333 s), Reward (3.333-7 s), ITI (7-17 s). No-lick was implied by all four outcomes encoded as zeros.

** Details on threshold-crossing time and alternative models included in ***Methods: Derivation of threshold and alternative decoding models***.

To exclude the sensory- and motor-related transients locked to the cue and the first-lick events in the threshold-crossing nests, the ramping interval was conservatively defined as 0.7 s post-cue up until 0.6 s before first-lick, and the minimum ramping interval for fitting was 0.1 s. Thus, for a trial to be included in the model, the first lick occurred between 1.4 s to 17 s (end of trial).

Initial model goodness of fit was assessed by R^2^, mean-squared loss and BIC. Models were 5-fold cross-validated with ridge regression at each nest to derive the final models, as described above. 95% confidence intervals on model coefficients were calculated by 2-sided t-test with standard errors propagated across sessions.

## Derivation of threshold and alternative decoding models

### Derivation of threshold models

As a metric of the predictive power of ramping DAN signals on first-lick time, we derived a threshold-crossing model. A threshold-crossing event was defined as the first time after the cue when the photometry signal exceeded and remained above a threshold level up until the time of first-lick on each trial. Importantly, while the analysis approach is reminiscent of pacemaker-accumulator models for timing, we make <u>no</u> claims that the analysis is evidence for pacemaker-accumulator models. Rather threshold-crossing times provided a convenient metric to compare the rate of increase in signals between trials.

Photometry timeseries for GCaMP6f and tdt were de-noised by smoothing with a 100 ms Gaussian kernel (kernel was optimized by grid screen of kernels ranging between 0-200 ms to minimize noise without signal distortion). To completely exclude the sensory- and motor-related transients locked to the cue and the first-lick events, the ramping interval was conservatively defined as 0.7 s post-cue up until 0.6 s before the first-lick. To eliminate chance crossings due to noise, we imposed a stiff, debounced threshold condition: to be considered a threshold crossing event, the photometry signal had to cross the threshold from low-to-high and remain above this level until the end of the ramping interval.

To derive an unbiased threshold for each session, we tested 100 evenly-spaced candidate threshold levels spanning the minimum-to-maximum photometry signal during the ramping interval for each session. Depending on threshold level, some trials never crossed, *i.e*., signal always remained below threshold or started and ended above threshold. Thus, the lowest candidate threshold for which there was a maximum number of trials crossing during the timing interval was selected as the “mid-level” threshold-crossing point. This threshold was specific to each photometry signal tested on each session. Threshold-crossing time was included in the decoding model as the normalized time on the ramping interval (0,1). If a trial never crossed threshold, it was encoded as a zero. If no trials ever crossed threshold, the threshold predictor was encoded as a vector of ones, thus penalizing the model for an additional predictor but providing no new information.

### Multi-threshold Model

An alternative model employed 3 unbiased thresholds: 1) the lowest threshold with ≥50 trials crossing (“min”); 2) the lowest threshold with the most crossings (“mid,” described above); and 3) the highest threshold with ≥50 trials crossing (“max”). For tdt datasets, trials rarely met the monotonic threshold constraint (usually the signals oscillated above and below the threshold throughout the ramping interval, failing to meet the debouncing constraint). Thus, to include tdt signals as conservatively as possible, we relaxed the 50-trial minimum constraint, taking the threshold with the most trials crossing, which was usually around 10 or fewer. The addition of more thresholds did not substantially improve the cross-validated model compared to the single-threshold model (***Figure 6—figure supplement 1***).

### Principal component analysis (PCA) threshold-crossing models

In another version of the decoding model, the threshold-crossing procedures were applied to ramping intervals fit with the first three PCs (as described in ***Methods: Principal Component Analysis (PCA)***) to derive a PCA version of the single-threshold and multi-threshold models. PCA analysis on tdt datasets showed no consistent PCs, and thus these PCs were not included in the decoding model. Instead, the actual tdt data was employed in the threshold model as in the other models described.

## Hierarchical Bayesian Modeling of Single-trial Dynamics

The probability of each single-trial SNc GCaMP6f signal belonging to a ramp vs. step Model Class was determined via Hierarchical Bayesian Model fitting with probabilistic programs written in the novel probabilistic programming language, Gen.jl, which is embedded in the Julia Programming Language (***Cusumano-Towner et al., 2019***). The top of the model hierarchy was the model class (linear ramp *vs*. step function) and the lower level was the respective parameterization of the two model classes (described below).

The probability of the step *vs*. ramp model class was inferred with data-driven inference. The best fit (step or ramp and parameterization) for each trial was calculated across 20 iterations (Gen *Traces*) of hierarchical modeling with 50 rounds of probabilistic refinement (computation *via* Gen Importance Resampling) per iteration (in model testing, models typically converged to their steady-state probability of model class within only 30 rounds of refinement, but 50 rounds were used conservatively to reduce the likelihood of suboptimal classifications).

*<u>Data-driven inference procedure</u>:* Each iteration of model fitting began at the top level of the hierarchy with a coin toss: with 50% probability, the probabilistic program would initialize with a model of either the Ramp or Step class. For data-driven inference, a Gen *Proposal* for the parameterization for this model class was then probabilistically generated. Data-driven proposals were designed to improve fitting efficiency and reduce computation time, and this allowed for faster convergence and better model fits as determined by the fit log-likelihood. The proposal heuristics were as follows:

<u>Ramp model</u>: A data-driven proposal was generated by dynamic noise random sample consensus (RANSAC (***Cusumano-Towner and Mansinghka, 2018***)) with additional data-driven constraints (see function ransac_assisted_model_selection_proposal in the Gen Github files):

1. SLOPE, *a*. The maximum data-supported slope was used to set the variance of slope sampling:

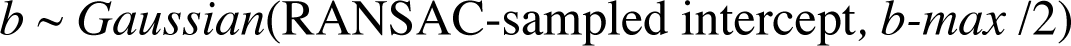

where *maxslope* was defined as the difference of the maximum and minimum signal within the trial dataset divided by the total duration of the trial (by definition, the largest slope supported by the data).
2. INTERCEPT, *b*. The initial search for the intercept (“*b-max*”) was calculated as the intercept for the calculated *maxslope* parameter), and this was used to set the noise level on sampling of the intercept parameter:

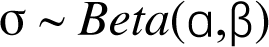
3. NOISE, σ. Parametrized noise level was sampled as:
where ɑ,β are the parameters of the beta distribution with mode=std(signal).

<u>Step model</u>: The data-driven proposal included two constraints/heuristics:

1. STEPTIME. *Derivative constraint*: To avoid sampling all unlikely step-times, *steptimes* were sampled uniformly from the timepoints where the derivative of the signal was in the highest 5% of the signal’s derivative across the trial dataset:
*steptime* ∼ *uniform*(indices of 95^th^ percentile of derivative of the signal)
2. LEFT and RIGHT SEGMENTS. Once a *steptime* was sampled, likely *left* and *right* segment amplitudes were sampled near the mean of the signal on either side of the step, *e.g.*:
*left ∼ Gaussian*(mean(signal left of *steptime*), std(signal left of *steptime*)) *right ∼ Gaussian*(mean(signal right of *steptime*), std(signal right of *steptime*))
3. NOISE, σ. The noise level was sampled as in the ramp model,

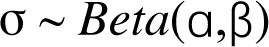

except ɑ,β were the parameterization of a Beta distribution with mode equal to the standard deviation of the signal left of *steptime*.

After model initialization for each Trace, 50 rounds of Importance Resampling of the hierarchical model were then conducted, each time randomly generating ramp or step hypotheses from the proposal heuristics. On each round, the best fitting hypothesis was retained, such that each of the 20 Trace iterations of model classification returned one optimized model from the 50 rounds of Importance Resampling.

The probability of the model class for each single-trial was then defined as the proportion of the 20 Trace iterations that found the optimal model to be derived from that model class (*e.g.,* if the model returned 15 step-fits and 5 ramp-fits, the p(ramp) was 0.25). Examples of the 20 Trace iterations for two sample trials are shown in ***Figure 6—figure supplement 2B*.**

To determine whether the step model detected step-functions in the GCaMP6f dataset, the step model was inferred alone to find step-fits for every trial, and single-trial signals were realigned to the optimal *steptime* (GCaMP6f, tdTomato, EMG, ***Figure 6—figure supplement 4A-B***).

## Single-trial dynamics analysis with geometric modeling (“Multiple threshold modeling”)

The multi-threshold procedure described above was also employed to determine whether single-trial ramping dynamics were more consistent with a continuous ramp *vs*. discrete step dynamic on single-trials. The threshold-crossing time for each trial was regressed against its first-lick time, and the slope of this relationship was reported, as well as the variance explained.

## Single-trial variance analysis for discrete step dynamics

For discrete step single trial dynamics to produce ramping on average, the time of the step across trials must be distributed throughout the trial interval (importantly, a peri-motor spike occurring consistently just before first-lick *cannot* give rise to continuous ramping dynamics on average). As such, the variance in the GCaMP6f signals across trials for similar first-lick times should be minimal near the time of the cue (when few trials have stepped) and near the time of the first-lick (when all of the trials have stepped). This predicts an inverted-U shaped relationship of signal variance across trials *vs*. position in the timing interval.

To compare variance across trials equitably, trials were first aligned to the cue and pooled by first-lick time in pools of 1s each (1-2 s, 2-3 s, *etc*.) truncating at the earliest first-lick time within the pool. The variance in GCaMP6f signals across trials within a pool was quantified in 10% percent increments of time from the cue up to the earliest first-lick time in the pool (*i.e*, 1-2 s pool truncated at 1 s, divided into 100 ms increments). Measuring variance by percent of elapsed time within pool allowed pooling of trials across the entire session. The shape of the variance *vs*. percent of timed interval elapsed was compared to the inverted-U shape prediction to assess for discrete step dynamics.

## Optogenetics—determining the physiological range for activation experiments

To test whether optogenetic manipulations during the self-timing task were in the physiological range, we assessed the magnitude of the effect of activation on dopamine release in the DLS by simultaneous photometry recordings with optical activation (***Figure 7—figure supplement 2***). In two DAT-cre mice, we expressed ChrimsonR bilaterally in SNc DANs and the fluorescent dopamine indicator dLight1.1 bilaterally in DLS neurons. SNc cell bodies were illuminated bilaterally (ChrimsonR 550 nm lime or 660 nm crimson, 0.5-5 mW) on 30% of trials (10 Hz, 10 or 20 ms up-time starting at cue onset and terminating at first-lick). dLight1.1 was recorded with 35 µW 475 nm blue LED light at DLS. To avoid cross-talk between the stimulation LED and the photometry recording site, the brief stimulation up-times were omitted from the photometry signal and the missing points filled by interpolation between the adjacent timepoints.

In a few preliminary sessions, we also explored whether we could evoke short-latency licking (*i.e*., within a few hundred ms of the stimulation) if light levels were increased above the physiological range for DAN signals. Rather than eliciting immediate licking, higher light levels produced bouts of rapid, nonpurposive limb and trunk movements throughout stimulation, and task execution was disrupted. The animals appeared to have difficulty coordinating the extension of the tongue to touch the lick spout. Simultaneous DLS dopamine detection showed large, sustained surges in dopamine release throughout the period of stimulation, with an average amplitude comparable to that of the reward transient (***Figure 7—figure supplement 2, right***). This extent of dopamine release was never observed during unstimulated trials. Consequently, to avoid overstimulation in activation experiments, we kept light levels well below those that generated limb and trunk movements.

## Optogenetics—naïve/expert control sessions

To determine whether optogenetic stimulation directly elicited or prevented licking, licking behavior was first tested outside the context of the self-timed movement task on separate sessions in the same head-fixed arena but with no cues or behavioral task. Opsin-expressing mice were tested before any exposure to the self-timed movement task (“Naïve”) as well as after the last day of behavioral recording (“Expert”). In ChR2 control sessions, stimulation (5 mW 425 nm light, 3 s duration, 10 Hz, 20% duty cycle) was applied randomly at the same pace as in the self-timed movement task. stGtACR2 control sessions were conducted similarly (12 mW 425 mW light, 3 s duration, constant illumination); but to examine if inhibition could block ongoing licking, we increased the baseline lick-rate by delivering juice rewards randomly (5% probability checked once every 5 s).

## Optogenetics—self-timed movement task

SNc DANs were optogenetically manipulated in the context of the 3.3 s self-timed movement task. To avoid overstimulation, light levels were adjusted to be subthreshold for eliciting overt movements as described above, and mice were not stimulated on consecutive days.

<u>Activation</u>: SNc cell bodies were illuminated bilaterally (ChR2: 0.5-5 mW 425 nm blue LED light;

ChrimsonR 550 nm lime or 660 nm crimson) on 30% of trials (10 Hz, 10 or 20% duty cycle starting at cue onset and terminating at first-lick). DAN terminals in DLS were stimulated bilaterally via tapered fiber optics on separate sessions.

<u>Inactivation</u>: SNc cell bodies were illuminated bilaterally (stGtACR2: 12 mW 425 nm blue light) on 30% of trials (constant illumination starting at cue onset and terminating at first lick).

## Quantification of optogenetic effects

The difference in the distribution of trial outcomes between stimulated and unstimulated trials on *each session* was quantified in four ways.

1. 2-Sample Unsigned Kolmogorov-Smirnov Test.
2. Difference in empirical continuous probability distribution function (cdf). The difference in the integral of the stimulated and unstimulated cdf (dAUC) was calculated for each session from 0.7-7 s. Effect size was quantified by permutation test, wherein the identity of each trial (stimulated or unstimulated) was shuffled, and the distribution of dAUCs for the permuted cdfs was calculated 10,000x. Results were reported for all sessions.
3. <u>3. Difference in mean movement time</u>. Movement times on stimulated and unstimulated trials were pooled and the distribution of movement time differences was determined by non-parametric bootstrap, in which a random stimulated and unstimulated trial were drawn from their respective pools 1,000,000x and the difference taken. The mean of each session’s bootstrapped distribution was compared across sessions by the 1,000,000x bootstrapped difference of the mean between sessions of different categories.
4. Difference in median movement time. Same as above but with median.

## Single-trial probabilistic movement state decoding model

The probability of transitioning to a movement state, s_t_=1, at time=t was decoded with a logistic generalized linear model of the form:

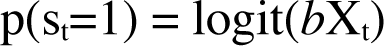

where X_t_ is a vector of predictors for the timepoint, *t*, and *b* is the vector of fit coefficients. The vector of predictors was comprised of the GCaMP6f signal at every timepoint (the current time, *t*) as well as the signal history, represented as 200 ms-wide signal averages moving back in time from *t*. Previous trial history (n-1^th^ and n-2^th^ first-lick times and reward/no-reward outcomes) did not contribute significantly to the model during model selection and were thus omitted (see Model Selection, below).

Movement state, s_t_, was defined as a binary variable, where state=0 represented all timepoints between the cue up until 160 ms before the first-lick detection (to exclude any potential peri-movement responses), and state=1 represented the timepoint 150 ms before the first-lick. Because there were many more state=0 than state=1 samples in a session, state=0 points were randomly down-sampled such that states were represented equally in the fit. To avoid randomly sampling a particular model fit by chance, each dataset was fit on 100 randomly down-sampled (bootstrapped) sets, and the average fit across these 100 sets was taken as the model fit for the session.

GCaMP6f signals were smoothed with a 100 ms gaussian kernel and down-sampled to 100 Hz. The GCaMP6f predictors were then nested into the model starting with those furthest in time from the current timepoint, *t*:

Nest 0: b0 (Null model)
Nest 1: b1 = b0 + mean GCaMP6f 1.8:2.0 s before current time=*t*
Nest 2: b2 = b1 + mean GCaMP6f 1.6:1.79 s before current time=*t*
Nest 3: b3 = b2 + mean GCaMP6f 1.4:1.59 s before current time=*t*
Nest 4: b4 = b3 + mean GCaMP6f 1.2:1.39 s before current time=*t*
Nest 5: b5 = b4 + mean GCaMP6f 1.0:1.19 s before current time=*t*
Nest 6: b6 = b5 + mean GCaMP6f 0.8:0.99 s before current time=*t*
Nest 7: b7 = b6 + mean GCaMP6f 0.6:0.79 s before current time=*t*
Nest 8: b8 = b7 + mean GCaMP6f 0.4:0.59 s before current time=*t*
Nest 9: b9 = b8 + mean GCaMP6f 0.2:0.39 s before current time=*t*
Nest 10: b10 = b9 + GCaMP6f signal at current time=*t*

Nesting the predictors from most distant in time to most recent permitted observation of the ability of more proximal signal levels to absorb the variance contributed by more distant signal history.

The fitted hazard function was then found as the average probability of being in the movement state across all trials in the session as calculated from the average model fit. Because s_t_=0 states were significantly down-sampled during fitting, this rescaled the fit hazard. Thus, to return the fit hazard to the scale of the hazard function calculated from the behavioral distribution, both the fit hazard and true hazard function were normalized on the interval (0,1), and the goodness of fit was assessed by R^2^ comparison of the fit and true hazard functions. This metric was similar between individual session fits as well as the grand-average fit across all animals and sessions.

To guard against overfitting, this procedure was repeated on the same datasets, except the datasets were shuffled before fitting to erase any non-chance correlations between the predictors and the predicted probability of being in the movement state.

## Model selection

To evaluate the contribution of task performance history to the probability of being in the movement state at time=*t*, we could not observe every timepoint in the GCaMP6f trial period timeseries as we did in the final model because the trial history for a given timepoint was the same for all other points in the trial; hence this created bias because the movement state=1 was represented for all trials, but the likelihood of the a trial’s 0 state being represented after down-sampling was dependent on the duration of the trial (*i.e.,* first-lick time). Consequently, model selection was executed on a modified version of the model that ensured that each trial would only be represented *one time at most* in the fit. Because this greatly reduced the power of the model, model selection was conducted on sessions from the two animals with the highest S:N ratio and most trials to ensure the best chance of detecting effects of each predictor (***Figure 8—figure supplement 1***).

The set of permutations of GCaMP6f signal and task history were fit separately, and the best model selected by BIC (though notably AIC and AICc were in agreement with the BIC selection). Each model was fit in “time-slices”—windows of 500 ms from the time of the cue up until the first-lick. Only one point for each trial was fit within this window to ensure the movement state within the window was uniquely represented. For each time-slice model, the GCaMP6f signal for each trial was thus averaged within the time-slice window, and the movement state was 1 only if the movement state occurred sometime within the window. The model fit for a session was taken as the average model fit across each of the time-slices. Notably, a time-slice required a sufficient number of trials to be present (either in the s_t_=0 or terminating in the movement state s_t_=1) for the fit to converge; once the first-lick occurred for a trial, it did not contribute data to later time-slices.

The source data files for ***Figure 8—figure supplement 1*** contain plots of all time-slice coefficient fits, including for models with insufficient numbers of trials to converge.

## Code Availability

All custom behavioral software and analysis tools are available with sample datasets at https://github.com/harvardschoolofmouse.

## Data Availability

All datasets supporting the findings of this study are publicly available https://www.dropbox.com/sh/g671nzziba405e3/AAByufom8BYs7PctECCbdiPua?dl=0

(Repository to be moved to Zenodo for publication). Source data files have been provided for all figures.

## ACKNOWLEDGEMENTS

We thank D. Chicharro, J.G. Mikhael, S.J. Gershman, K. Reinhold, S. Panzeri, R. Bliss and E.N. Brown for discussions on analytical methods; J. Tenenbaum, C. Wong, and A. Lew for their instruction and advice pertaining to probabilistic programming in Gen; V. Berezovskii, J. LeBlanc, T. LaFratta, O. Mazor, P. Gorelik, J. Markowitz, A. Lutas, K.W. Huang, L. Hou, S.J. Lee and N.E. Ochandarena for technical assistance; and B. Sabatini, C. Harvey, S.R. Datta, M. Andermann, L. Tian and W. Regehr for reagents. The work was supported by NIH grants UF-NS108177 and U19-NS113201, and NIH core grant EY-12196. A.E.H. was supported by a Harvard Lefler Predoctoral Fellowship and a Stuart H.Q. & Victoria Quan Predoctoral Fellowship at Harvard Medical School. The funders had no role in study design, data collection and interpretation, or the decision to submit the work for publication.

## AUTHOR CONTRIBUTIONS

A.E.H. and J.A.A conceived the project. A.E.H. performed all experiments. G.S. assisted with experiments using tapered fiber optics; Y.H. assisted with optogenetic no-opsin control experiments. F.S. and Y.L. developed the dopamine sensor, DA_2m_. A.E.H. and J.A.A. analyzed the data and wrote the paper.

## SUPPLEMENTAL FIGURES

**Figure 1—figure supplement 1.**
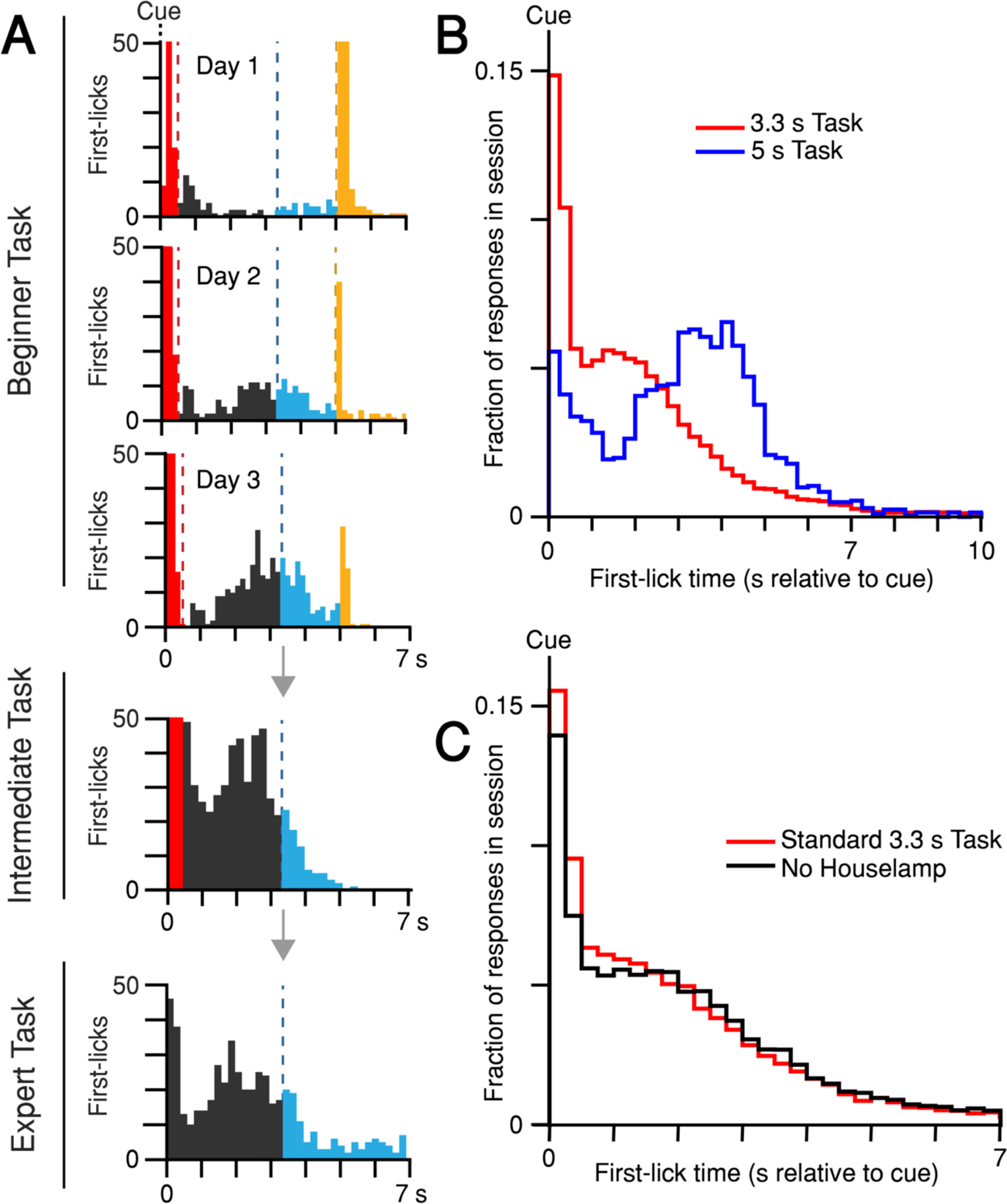
Self-timed movement task learning and variations. (**A**) Task learning. Histogram of first-lick times from single sessions at different stages of training (red: reaction, grey: early, blue: operant-rewarded, yellow: Pavlovian-rewarded). Bars >50 first-licks truncated for clarity. (**B**) Mice adjust behavior to the timing-contingencies of the task. First-lick time distributions from tasks with different target timing intervals. Red: 3.3 s reward-boundary. Blue: 5 s reward-boundary (all sessions, all mice). **(C)** Mice time their first-licks relative to the start cue, not the houselamp. First-lick time distributions during behavior with (red) and without (black) houselamp events (4 mice, 4-5 sessions/mouse on each version of the task). Source data: Figure 1***—source data***.

**Figure 1—figure supplement 2.**
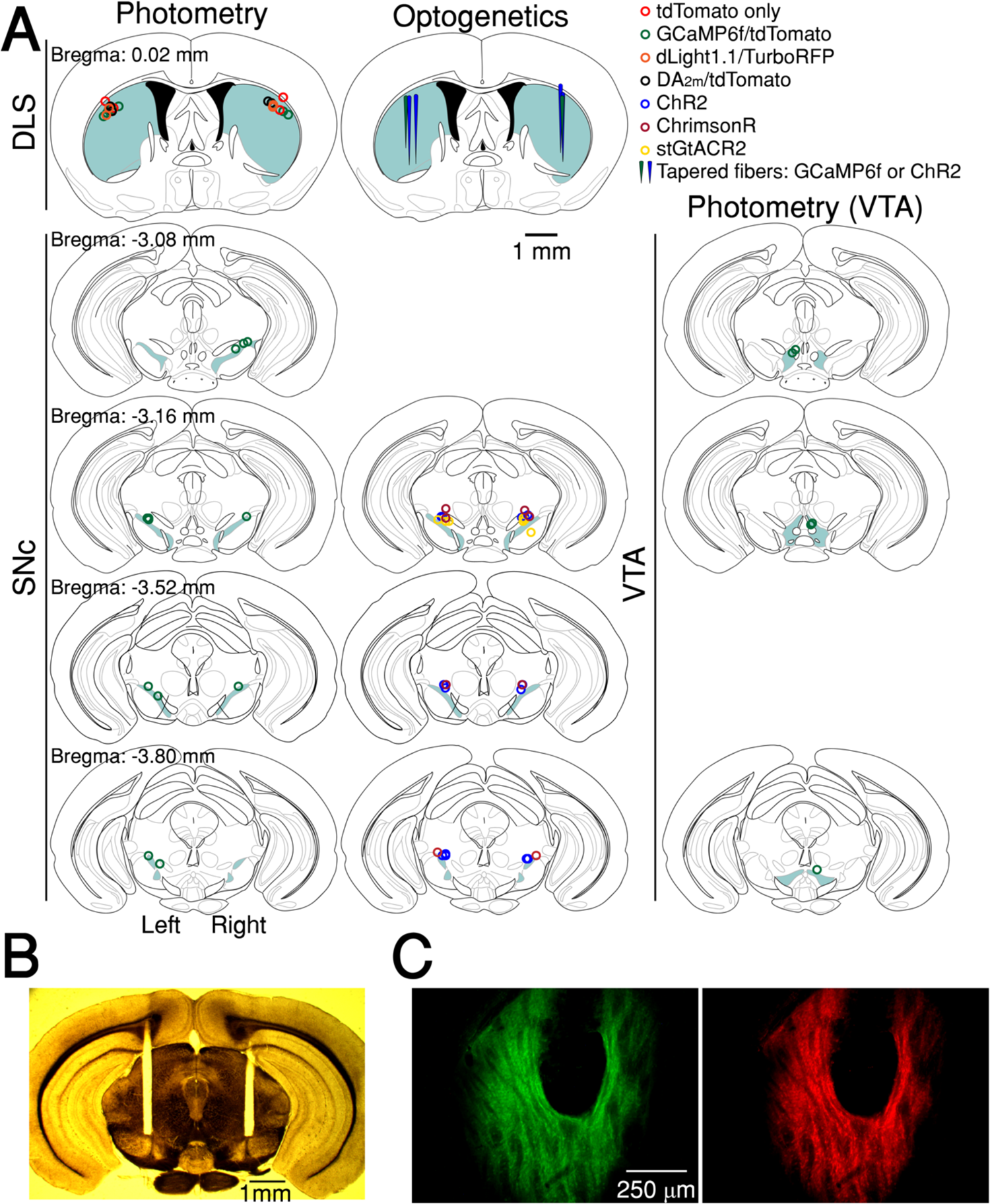
Fiber optic placement and histology**. (A)** Approximate fiber positions for all mice. **(B)** Brightfield microscopy with polarized filter on a freshly-cut brain slice showing bilateral fiber placement at SNc (from stGtACR2 experiment). **(C)** Example of co-expression of green (DA2m,) and red (tdTomato) fluorophores relative to fiber optic tip.

**Figure 2—figure supplement 1.**
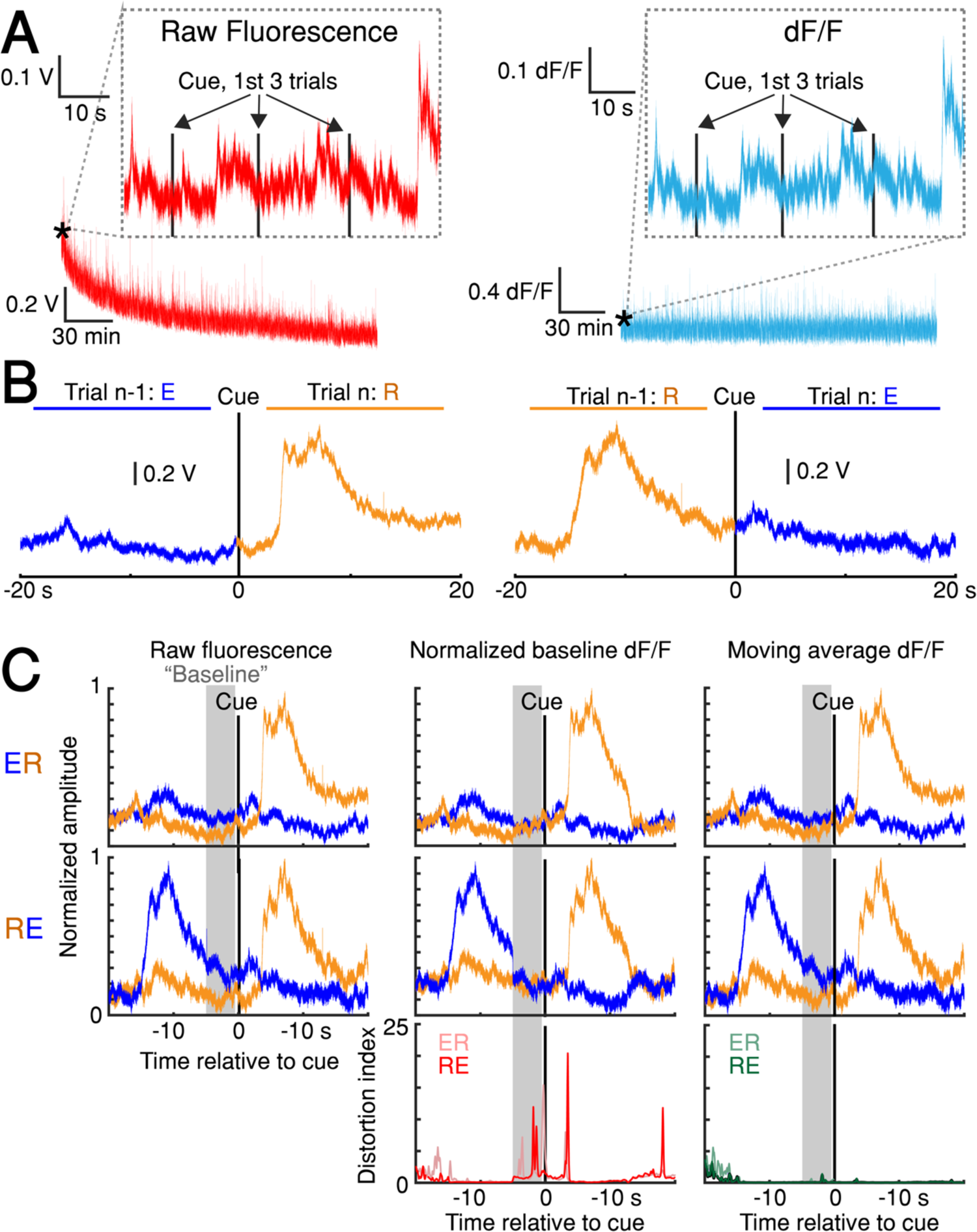
dF/F method validation**. (A)** Left: slow, raw fluorescence bleaching across one session. Left inset: Minimal bleaching occurs across the first 3 trials (∼1 min). Right: dF/F removes slow bleaching dynamics. Right inset: The same 3-trial window shown for dF/F signal. **(B)** Average raw fluorescence on paired, consecutive trials from one session aligned to cue on the n^th^ trial. Left: n-1^th^ trial was early, n^th^ trial was rewarded (“ER” condition). Right: “RE” condition (See ***<u>Methods: dF/F method characterization and validation</u>***<u>). **(C)** Comparison of baseline GCaMP6f signals on paired, consecutive trials aligned to cue. Columns: three different versions of the signal (Raw fluorescence, Normalized baseline dF/F method, Moving average dF/F method). Top row: ER condition; middle row: RE condition; bottom row: distortion index. Red distortion index plot shows only Normalized baseline method. Green distortion index plot shows overlay of Moving Average, Low-Pass Filter, and Multiple Baseline dF/F Methods because the difference in signal distortion between these methods was indistinguishable (See ***Methods: dF/F method characterization and validation***). Source data:</u> Figure 2***<u>—source data</u>***.

**Figure 2—figure supplement 2.**
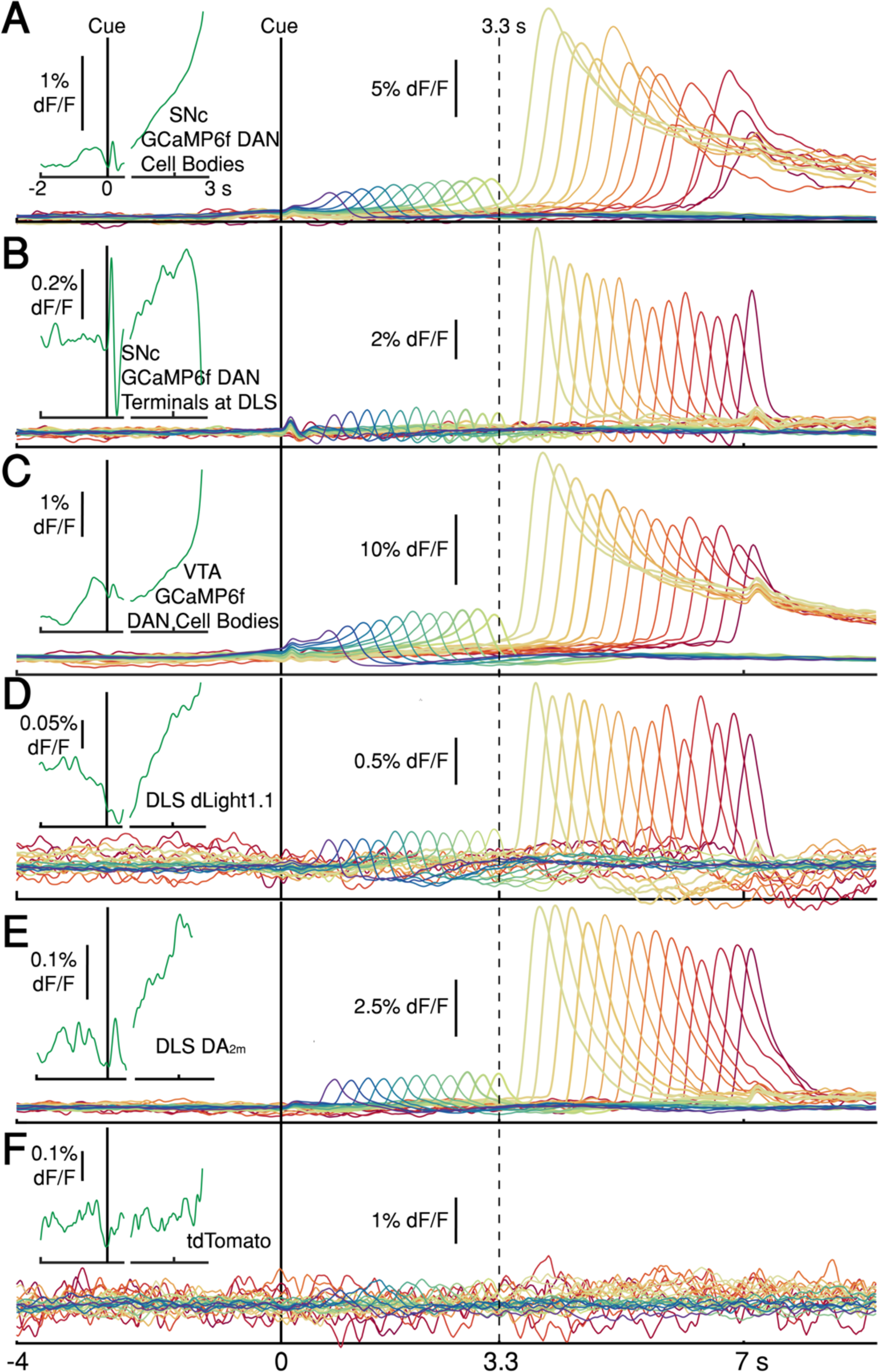
Average photometry signals, pooled every 250 ms by first-lick time, spanning 0.5 s (purple) to 7 s (red). Signals in main panels aligned only to cue, not first-lick. **(A)** Average DAN GCaMP6f signals at SNc cell bodies (12 mice). **(B)** DAN GCaMP6f signals at axon terminals in DLS (10 mice). **(C)** Striatal dopamine detection with dLight1.1 at DLS (5 mice). **(D)** Striatal DA_2m_ signals at DLS (4 mice). **(E)** DAN GCaMP6f signals at VTA cell bodies (4 mice). **(F)** tdTomato signals. **Insets (left)**: Cue and lick-aligned average signals for a single time bin before first-lick to show pre-lick ramping present in all dopaminergic signals. Left of axis break: aligned to cue. Right of axis break: aligned to first-lick. Traces plotted up until 150 ms before first-lick. Source data: Figure 2***—source data***.

**Figure 3—figure supplement 1.**
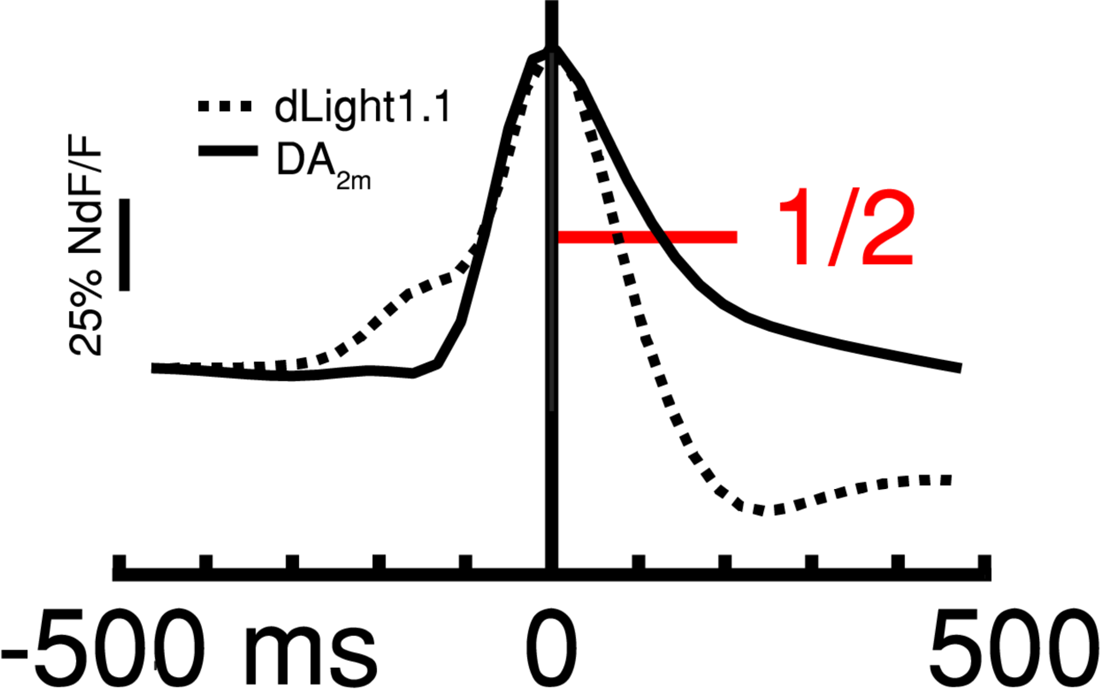
Comparison of dLight1.1 (dashed) and DA_2m_ (solid) kinetics surrounding peak of unrewarded transient (first-lick: 0.5-3.3 s). Red line: ½ baseline-to-peak amplitude for measuring decay t_1/2_ (see ***Methods***). Source data: Figure 3***—source data***.

**Figure 4—figure supplement 1.**
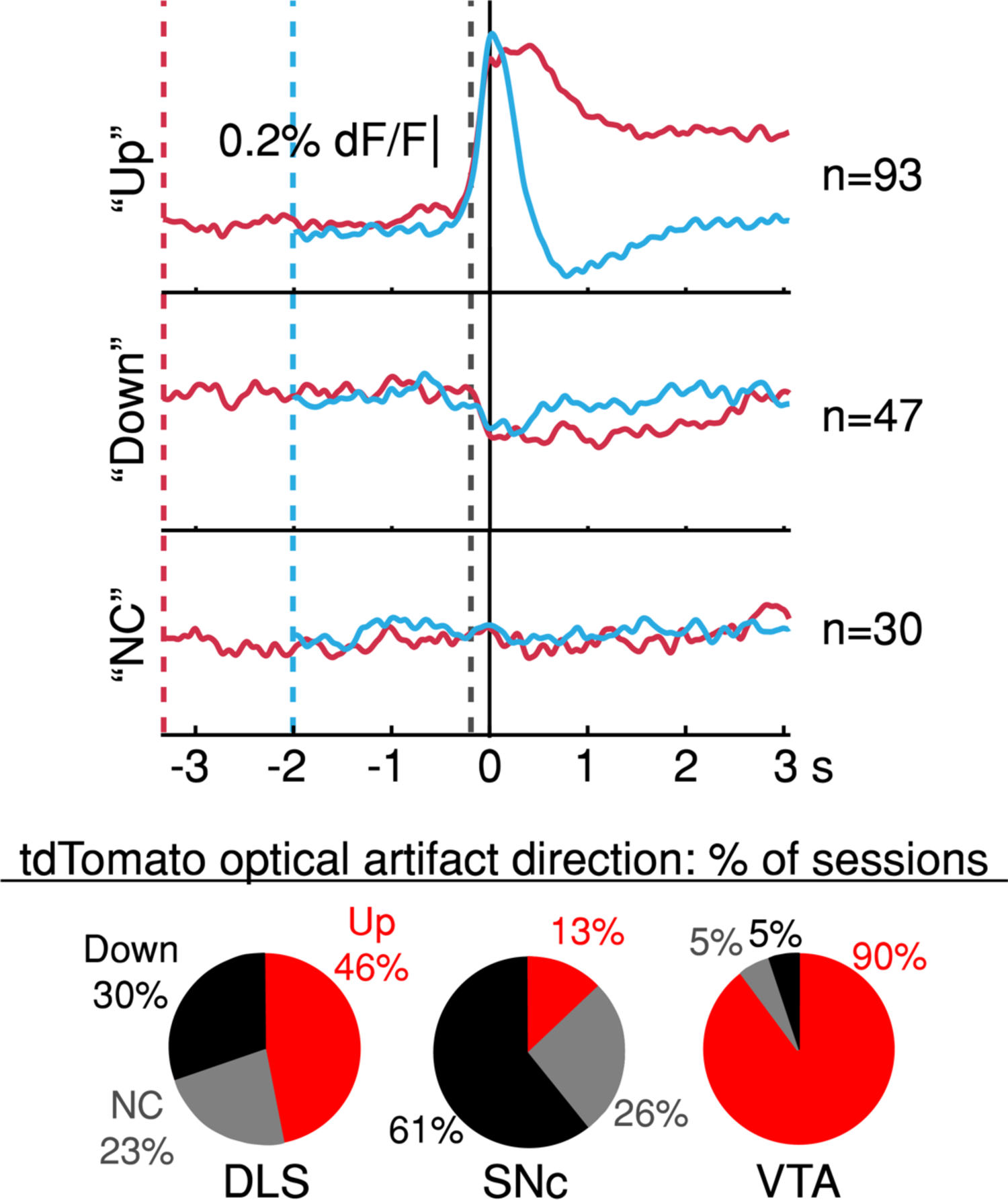
Average tdTomato optical artifacts (aligned to first-lick time) showed inconsistent directions even within the same session. Averages for all three types of artifact (consistently up, “Up”; consistently down, “Down”; and not consistent “NC”) shown for all sessions. Pie plots: Breakdown of average tdt artifact direction by session at each recording site. Source data: Figure 4***—source data***.

**Figure 5—figure supplement 1.**
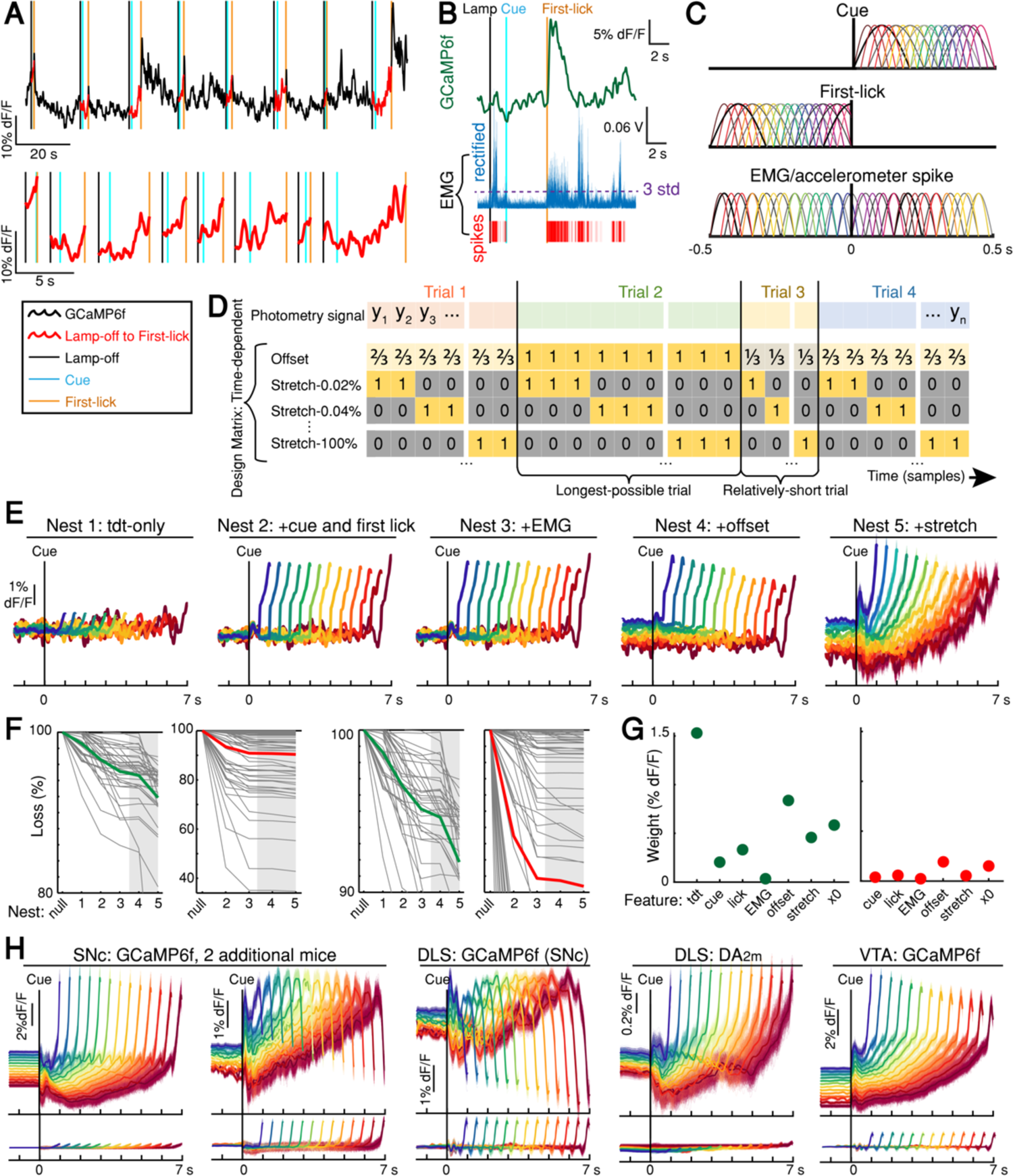
DAN signal encoding model parameterization and model selection. **(A)** Schematic of photometry timeseries fit by encoding model. The lamp-off to first-lick interval was excised from each trial in a session (top) and concatenated to produce the timeseries fit by the model (bottom). **(B)** EMG spikes derivation: thresholding rectified EMG at 3 standard deviations (example trial). **(C)** Optimized basis kernels to produce timing-independent features. **(D)** Schematic of Design Matrix for timing-dependent features. **(E)** GCaMP6f model fits by nest iteration for example session. Shading: model error simulated 300x. **(F)** Model loss by nest iteration. Green: mean loss for SNc GCaMP6f; red: mean loss for tdTomato (tdt); grey lines: individual sessions; grey shading: timing-dependent nests. Left: full-scale view of all datasets. Right: mean GCaMP6f and tdt loss compared on same scale. **(G)** Summary of feature weights across SNc GCaMP6f (left) and tdt (right) models. Coefficient weights were rectified, summed, and divided by the number of predictors per feature. 2x standard error bars (too small to see). All features were significant in both GCaMP6f and tdt models. **(H)** Top: examples of the full timing-dependent model (nest 5) from additional mice for all recorded dopaminergic signals. Bottom: tdt control channel fit. Model errors simulated 300x. Some mice show downward-going movement-related spikes at SNc cell bodies (second panel). All mice showed downward-going movement-related spikes from SNc terminals in DLS (middle panel). Source data: Figure 5***—source data***.

**Figure 5—figure supplement 2.**
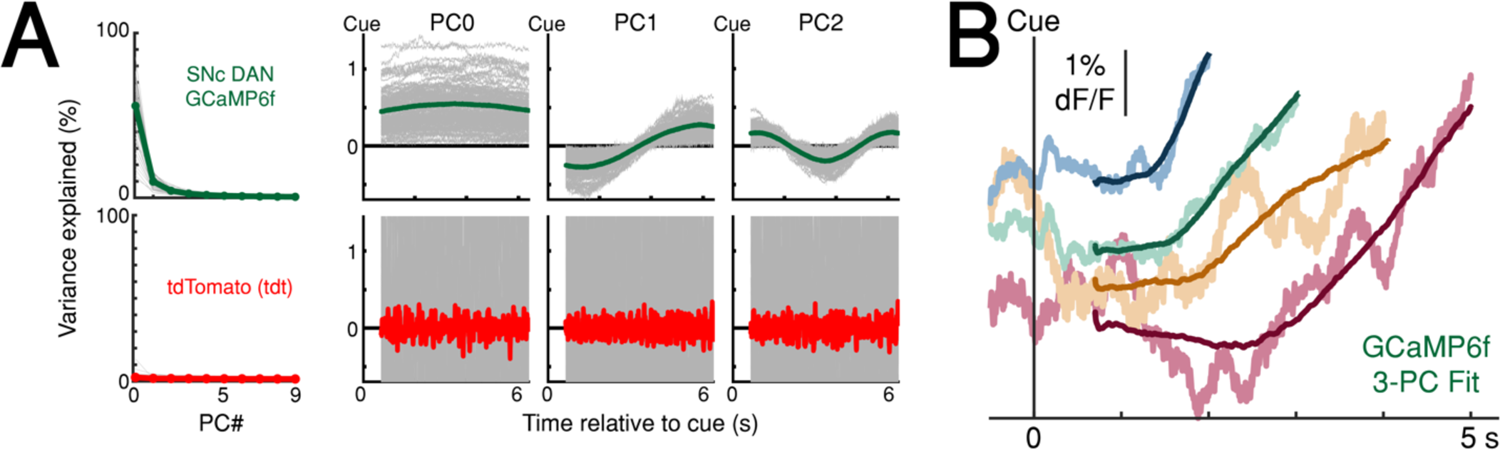
Principal component analysis (PCA) of the ramping interval (0.7 s up to first-lick relative to cue). **(A)** Left: Variance explained by first 10 principal components (PC). Right: first three principal components. Green line: mean PC, GCaMP6f recorded at SNc; Red line: mean PC, tdTomato (tdt) recorded at SNc and VTA; Grey lines: single-session data. X-axis shown for longest-possible interpolated trial duration; trials of shorter duration were interpolated to have the same number of samples for PCA. **(B)** Example session data simulated with first 3 PCs. Noisy traces: actual averaged GCaMP6f signals truncated at first-lick onset; Smooth traces: PC fits of the same trials. Source data: Figure 5***—source data***.

**Figure 6—figure supplement 1.**
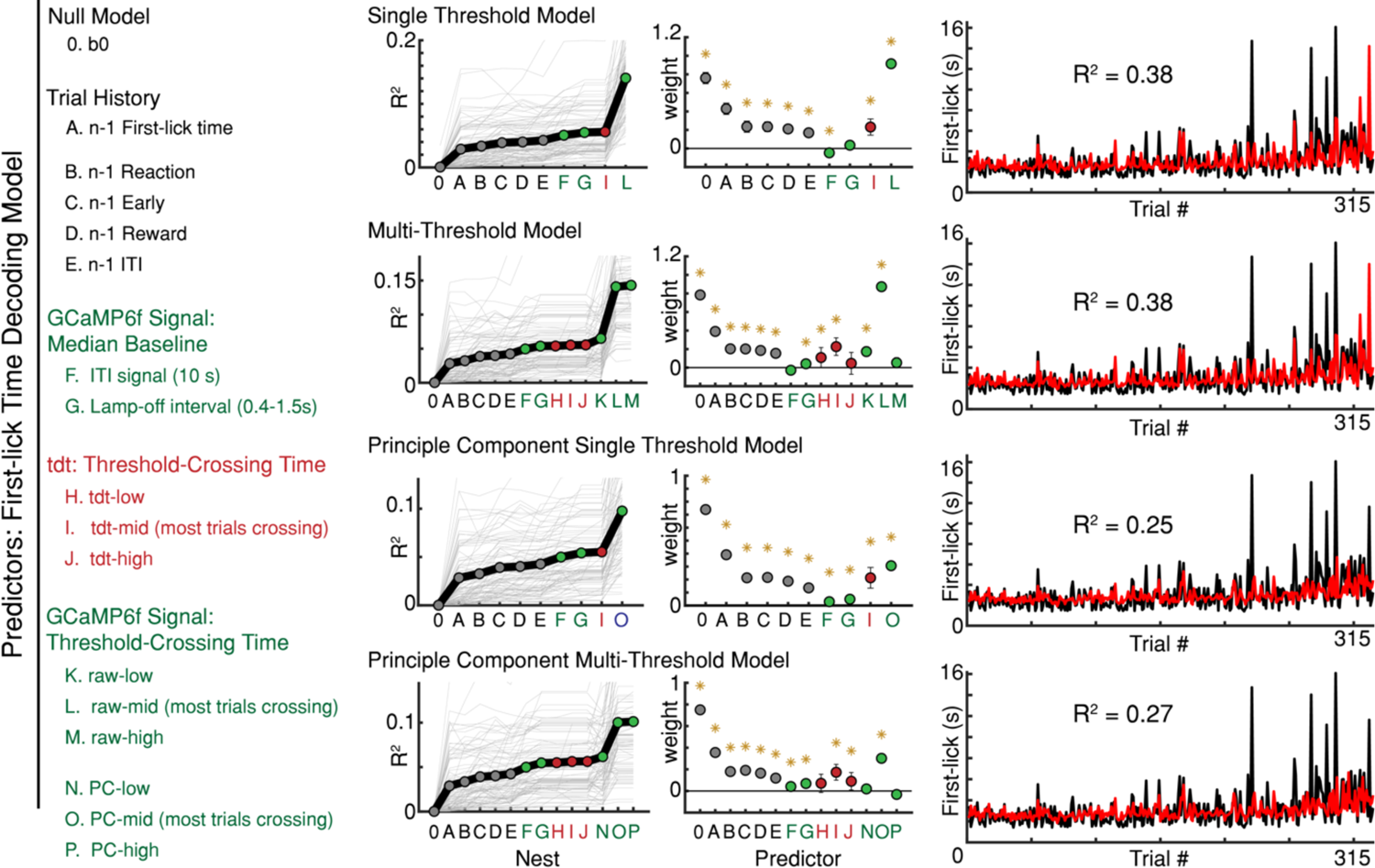
Variations of the first-lick time decoding model. *: p<0.05, error bars: 95% confidence intervals. GCaMP6f threshold crossing time dominated every version of the model; n-1^th^ trial first-lick time was consistently the second-best predictor. Source data: Figure 6***—source data***.

**Figure 6—figure supplement 2.**
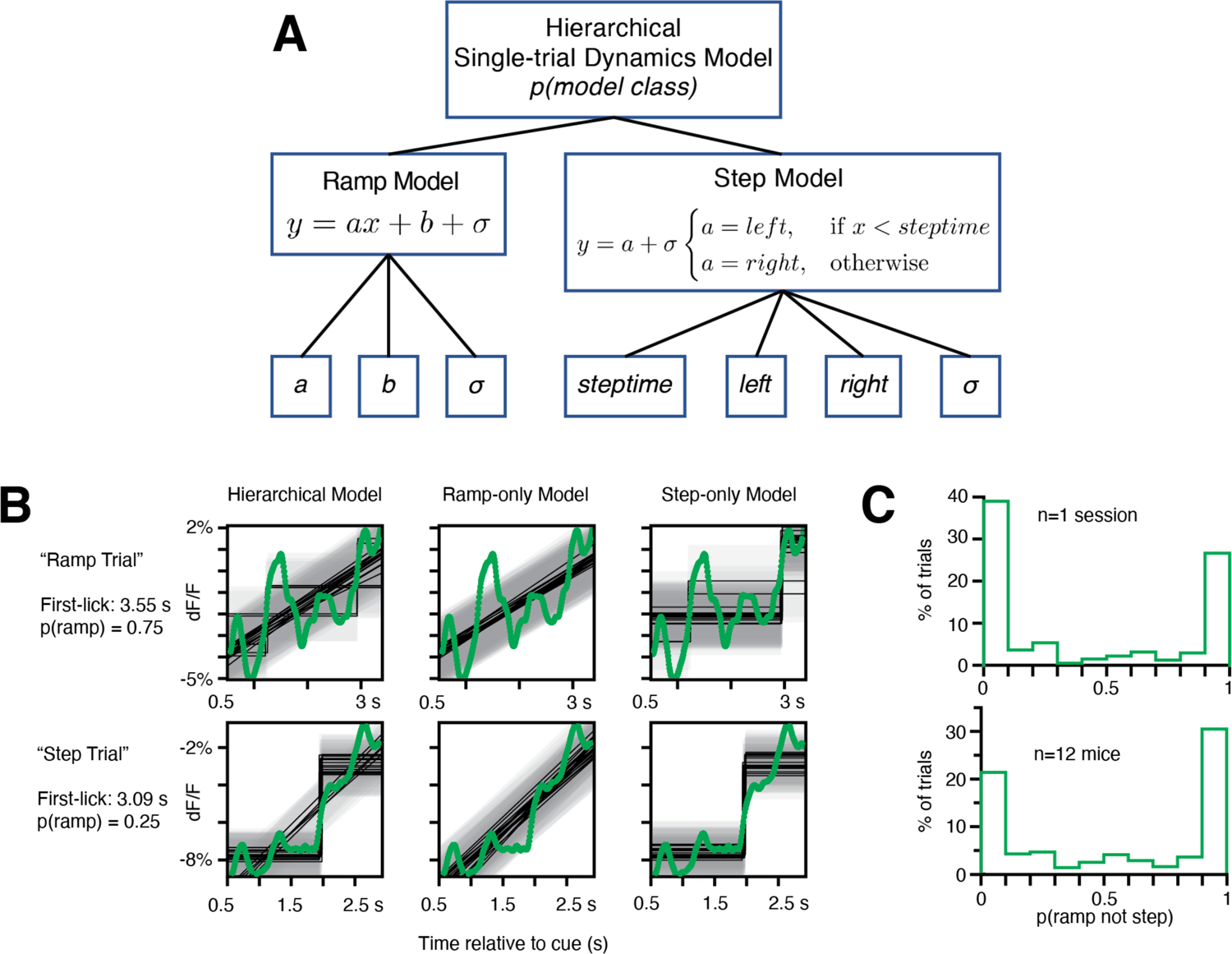
Analysis of single-trial dynamics: Hierarchical Bayesian Ramp *vs.* Step Modeling. **(A)** Schematic (see ***Methods: Hierarchical Bayesian Modeling of Single-trial Dynamics***). **(B)** Example fits from hierarchical model on 2 example single trials from the same epoch in a single session. Green: SNc GCaMP6f single-trial signal, light grey shading: noise band, dark grey lines: model fits. Note that the top trial is more frequently classified as a ramp, and the lower trial is more frequently classified as a step. However, both the ramp and step models return intuitive and reasonable fits to both single-trial signals. **(C)**. Probability of model class across all trials. X axis: 0 indicates all probabilistic fits for a given trial returned step-class models; 1 indicates all ramp-class models. Single sessions across mice showed considerable uncertainty in model classification. Source data: Figure 6***—source data***.

**Figure 6—figure supplement 3.**
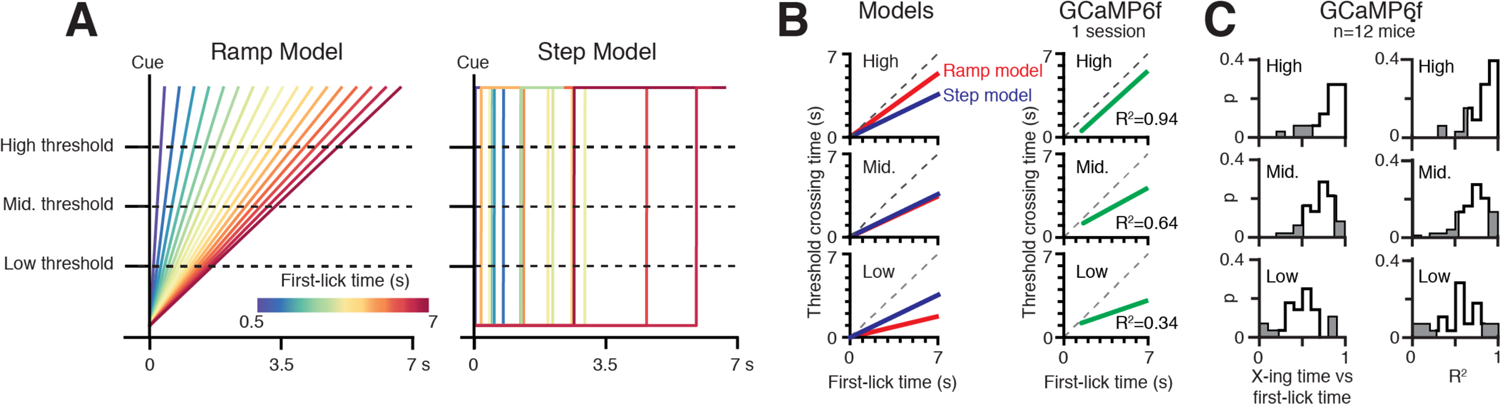
Geometric analysis of single-trial dynamics with Multiple Threshold Modeling. **(A)** Left: linear ramp model, Right: discrete step model. Step positions drawn from uniform distribution over the cue-to-first-lick interval. Low-, Mid- and High-level thresholds shown. **(B)** Threshold-crossing time *vs.* first-lick time (“X-ing time *vs*. first-lick time”) for (from top to bottom) High-, Mid- and Low-level thresholds. Left: simulation predictions for ramp and step models. Right: X-ing time *vs*. first-lick time regression fit on single trials from 1 session (data from Figure 6A). The step model predicts X-ing time *vs*. first-lick time does not change across threshold levels, whereas ramp model predicts the slope of this relationship increases as threshold is raised. Single-trial GCaMP6f data exhibits increasing X-ing time *vs*. first-lick time slope with increasing threshold level, consistent with the ramp model but inconsistent with the step model. **(C)**. X-ing time *vs*. first-lick time across all mice. Left column: frequency of slope relationship across sessions, right column: variance explained. Source data: Figure 6***—source data***.

**Figure 6—figure supplement 4.**
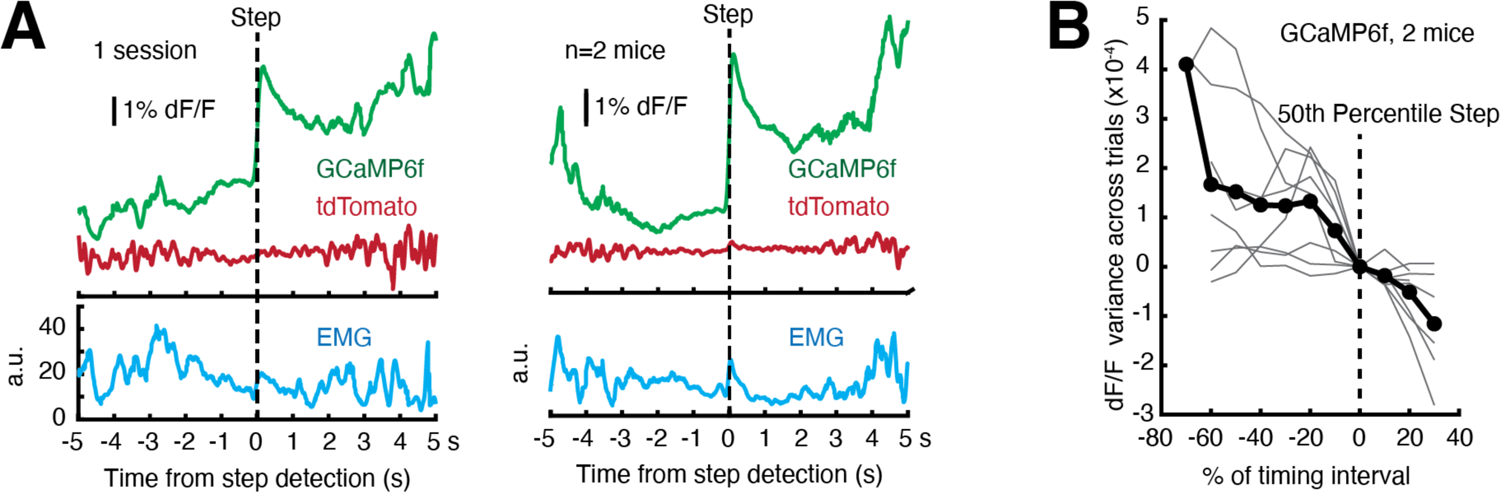
Assessing single-trial dynamics. **(A)** Single-trial signals aligned to discrete step position as found by Bayesian step model do not exhibit discrete step dynamics. To best estimate step times, the two animals with the highest GCaMP6f S:N were examined (Mouse B5 and B6). Left: 1 session, Right: average of signals from both mice. **(B)** Variance of GCaMP6f signals across trials. Step times were computed by Bayesian step model. An ideal step model predicts maximal variance at the 50^th^ percentile step, but variance declined monotonically on average. Grey lines: single sessions; black line: average. For detailed explanation, see ***Methods: Single-trial variance analysis for discrete step dynamics***. Source data: Figure 6***—source data***.

**Figure 7—figure supplement 1.**
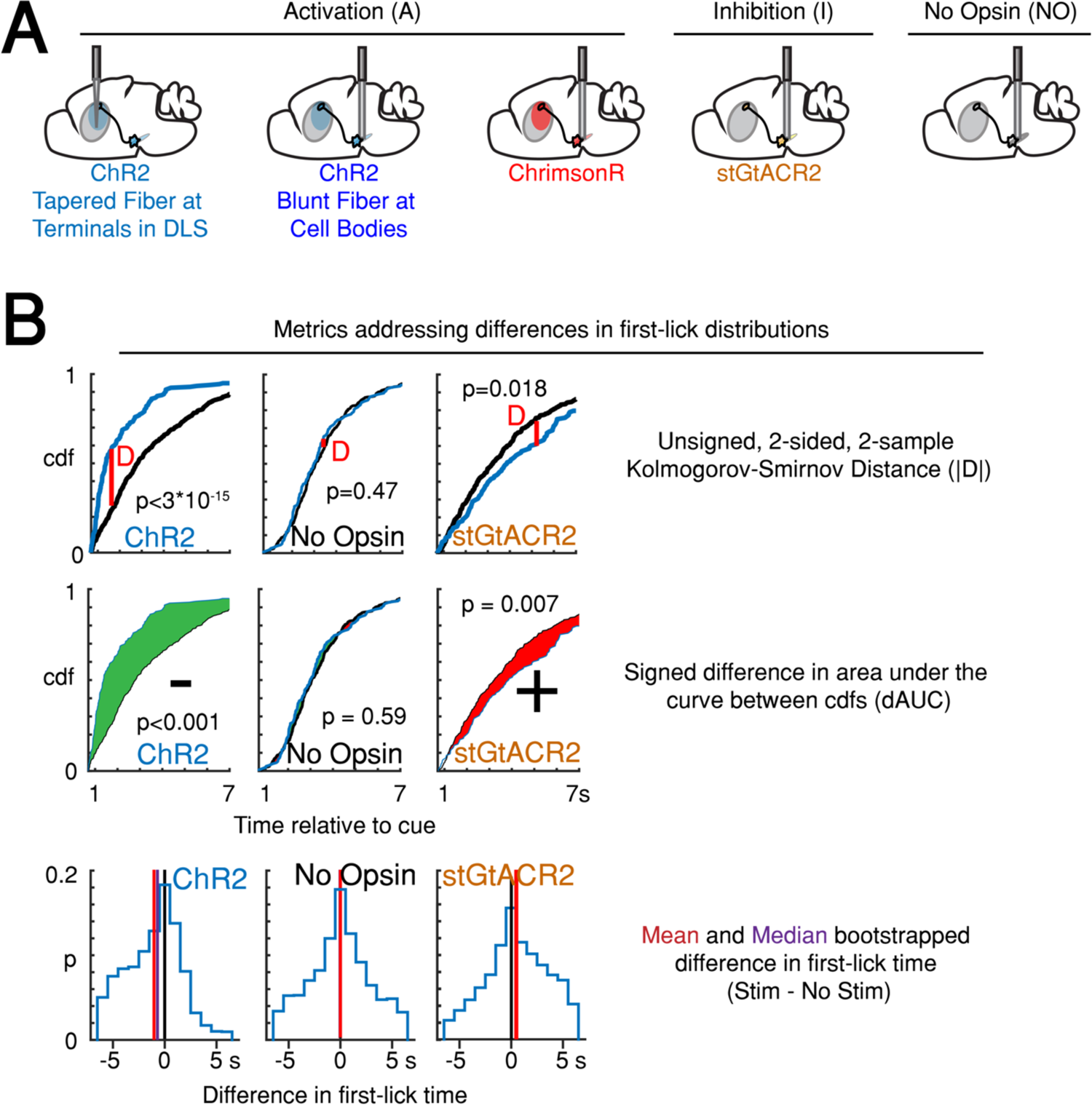
Variations on measurements of optogenetic effects. **(A)** Strategy for optogenetic targeting of DANs. **(B)** Comparison of four complementary metrics for addressing optogenetic effects. Left: unsigned Kolmogorov-Smirnov Distance (KS-D) analysis of differences in first-lick time distribution. Center: signed, bootstrapped comparison of difference in area under the cdf curves (dAUC). Right: mean and median bootstrapped difference in first-lick time. Source data: Figure 7***—source data***.

**Figure 7—figure supplement 2.**
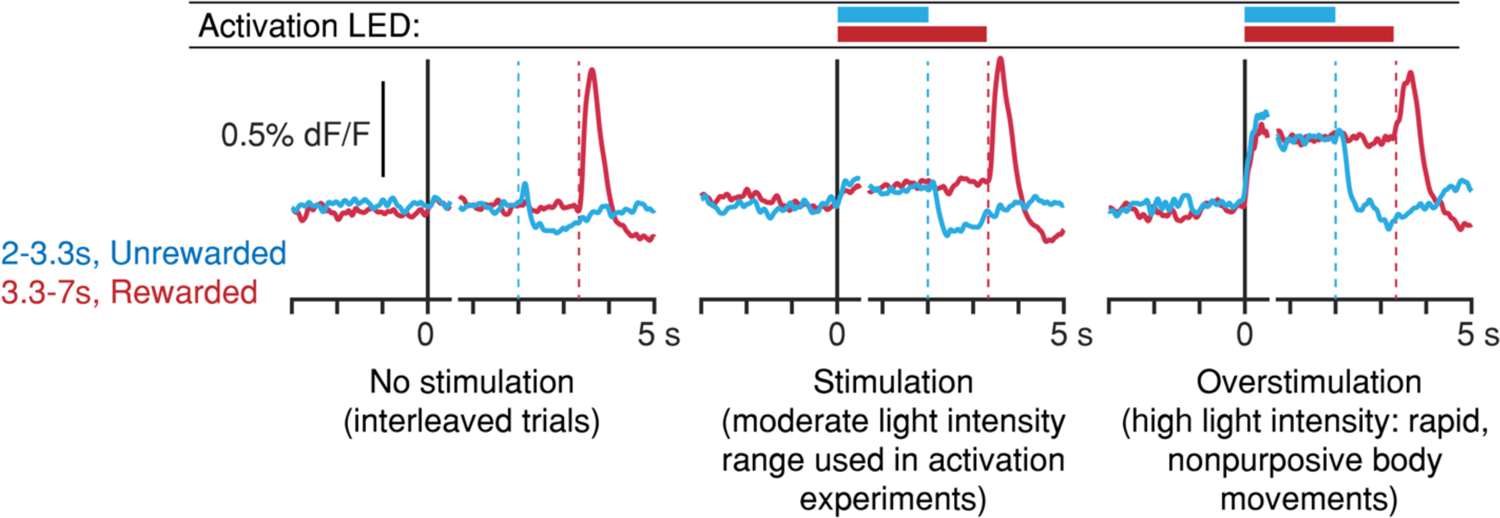
Light-power calibration for optogenetic activation of DANs. In preliminary experiments, DLS dopamine levels were monitored during the self-timed movement task, in which SNc DANs were activated randomly on 30% of interleaved trials. Dashed vertical lines: first-lick time. **Left**: interleaved, unstimulated trials (2 mice, 8 sessions). **Middle**: stimulated trials at the range of light levels used in the activation experiments show slightly elevated DLS dopamine signals compared to interleaved, unstimulated trials. First-lick timing was generally early-shifted in these sessions. **Right**: in a subset of preliminary calibration sessions, stimulation light levels were increased to the point where rapid, nonpurposive limb/trunk movements were observed throughout stimulation (1 mouse, 3 sessions). DLS dopamine signals show much higher, sustained increases throughout stimulation. Ongoing body movements disrupted task participation. Source data: Figure 7***—source data***.

**Figure 7—figure supplement 3.**
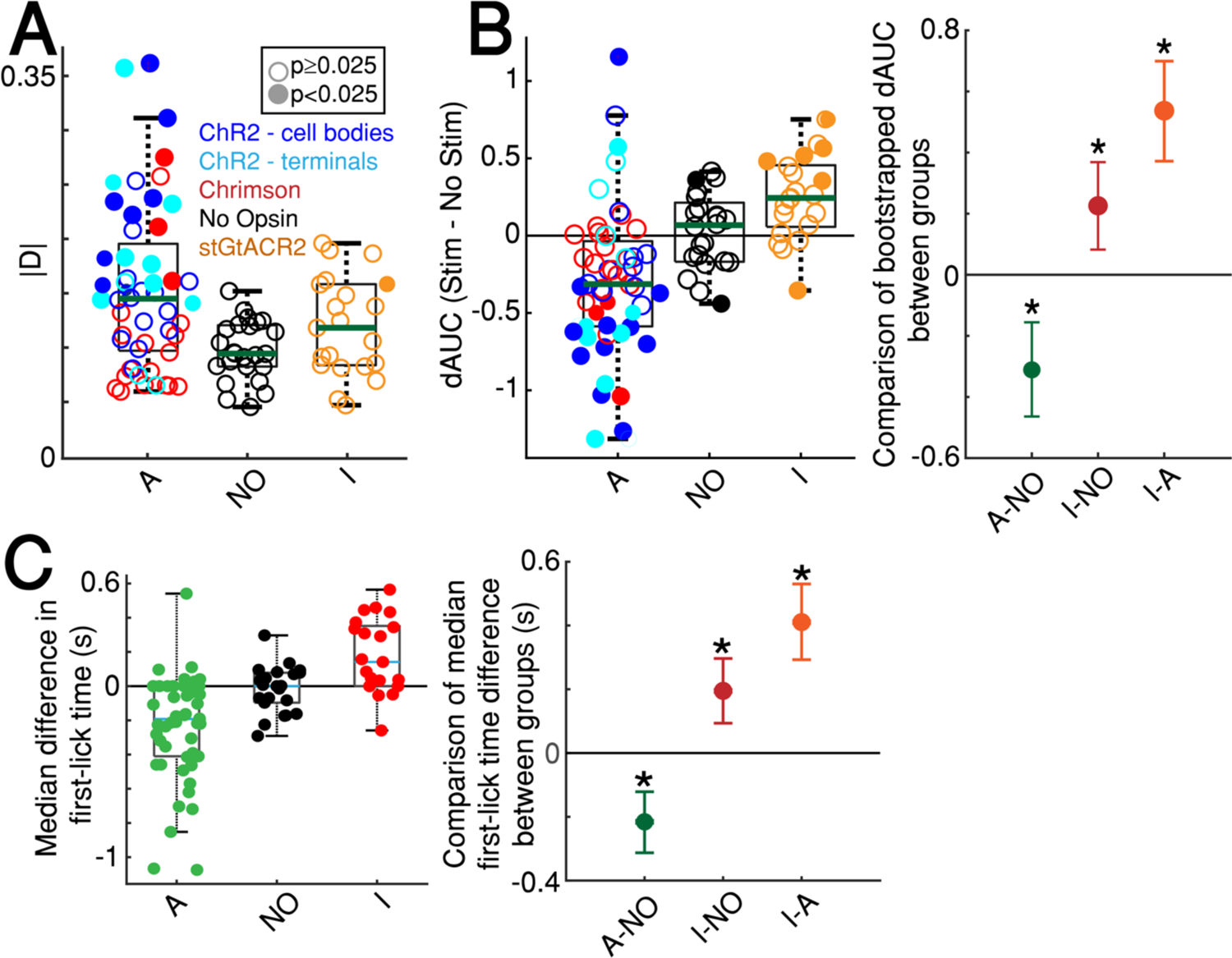
Quantification of optogenetic effects with additional metrics. **(A)** KS-D analysis: all sessions. “A”: activation sessions; “NO”: no opsin sessions; “I”: inhibition sessions. Filled circles indicate significant difference between stimulated/unstimulated trials on single session (p<0.025, 2-sided, 2-sample KS test). Standard box plot, line: median, box: upper/lower quartiles; whiskers: 1.5x IQR. **(B)** Left: bootstrapped dAUC Assay: all sessions, standard box plot as in *(**A**)*. Filled circles: significant difference on single session (p<0.025, 2-sided bootstrapped dAUC test, see ***Methods***). Right: comparison of dAUC in first-lick distributions across all sessions between groups. Error bars denote bootstrapped 95% confidence interval (*: p<0.05). **(C)** Left: median bootstrapped difference in first-lick time, stimulated-minus-unstimulated trials, standard box plot as in (***A***). Dots: single sessions. Right: Comparison of median difference in first-lick time across all sessions. Error bars denote bootstrapped 95% confidence interval (*: p < 0.05). Source data: Figure 7***—source data***.

**Figure 7—figure supplement 4.**
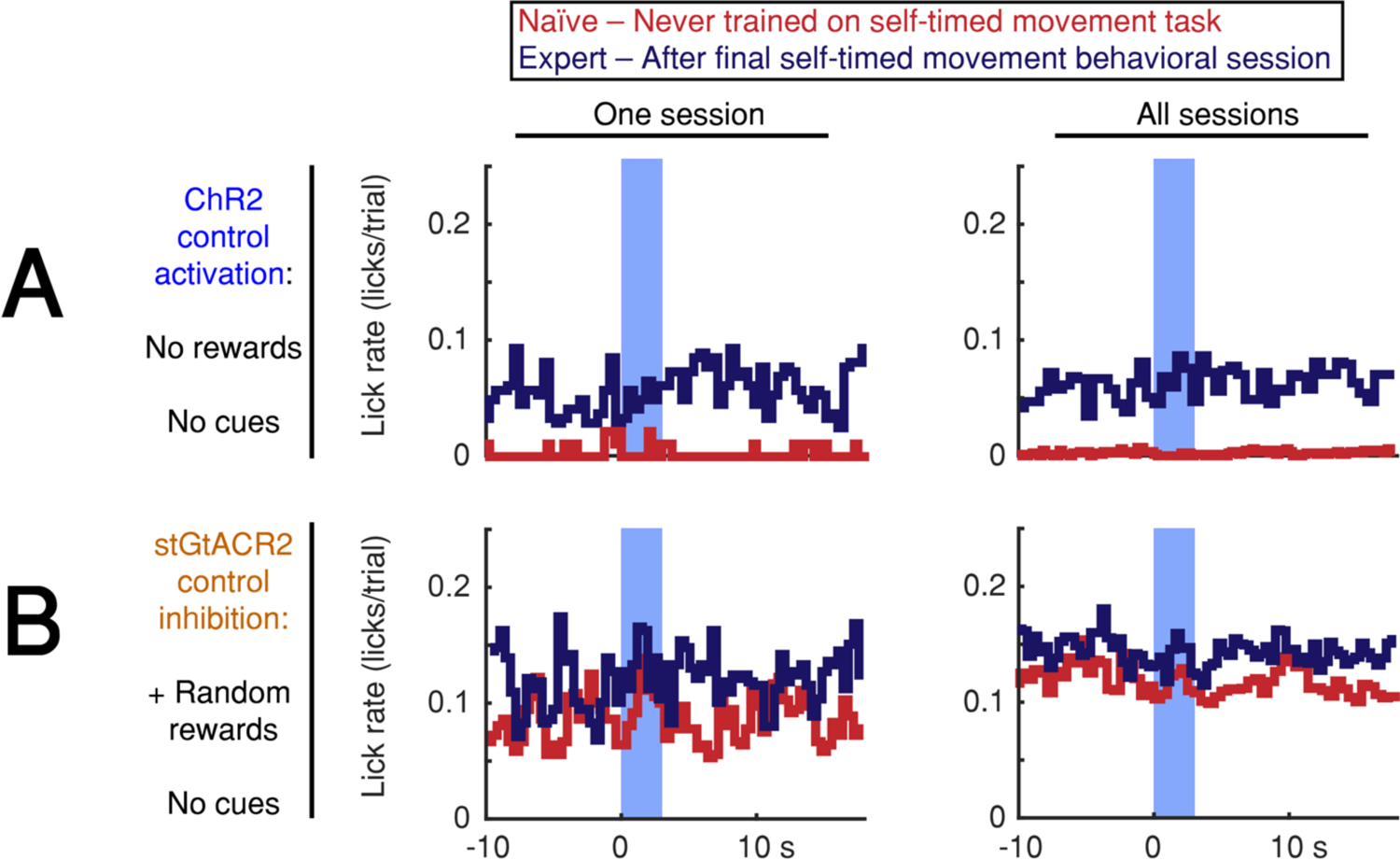
Optogenetic DAN stimulation does not cause or prevent licking. **(A,B)** Stimulation-aligned lick-rate during control sessions. Animals were tested in 1-3 control sessions both before exposure to the self-timed movement task (red) and in 1-2 control sessions after the end of behavioral training (navy). Blue bar indicates stimulation period (3 s). Left: one session, Right: all sessions. **(A)** Activation control sessions (no cues or rewards). Animals were head-fixed on the behavioral platform and stimulated randomly at the same pace as the standard 3.3 s self-timed movement task. Activation did not elicit immediate licking in any session. **(B)** Inhibition-control sessions (no cues, + random rewards). Animals were head-fixed on the behavioral platform while receiving juice rewards at random times. Inhibition did not prevent licking in any session. Source data: Figure 7***—source data***.

**Figure 8—figure supplement 1.**
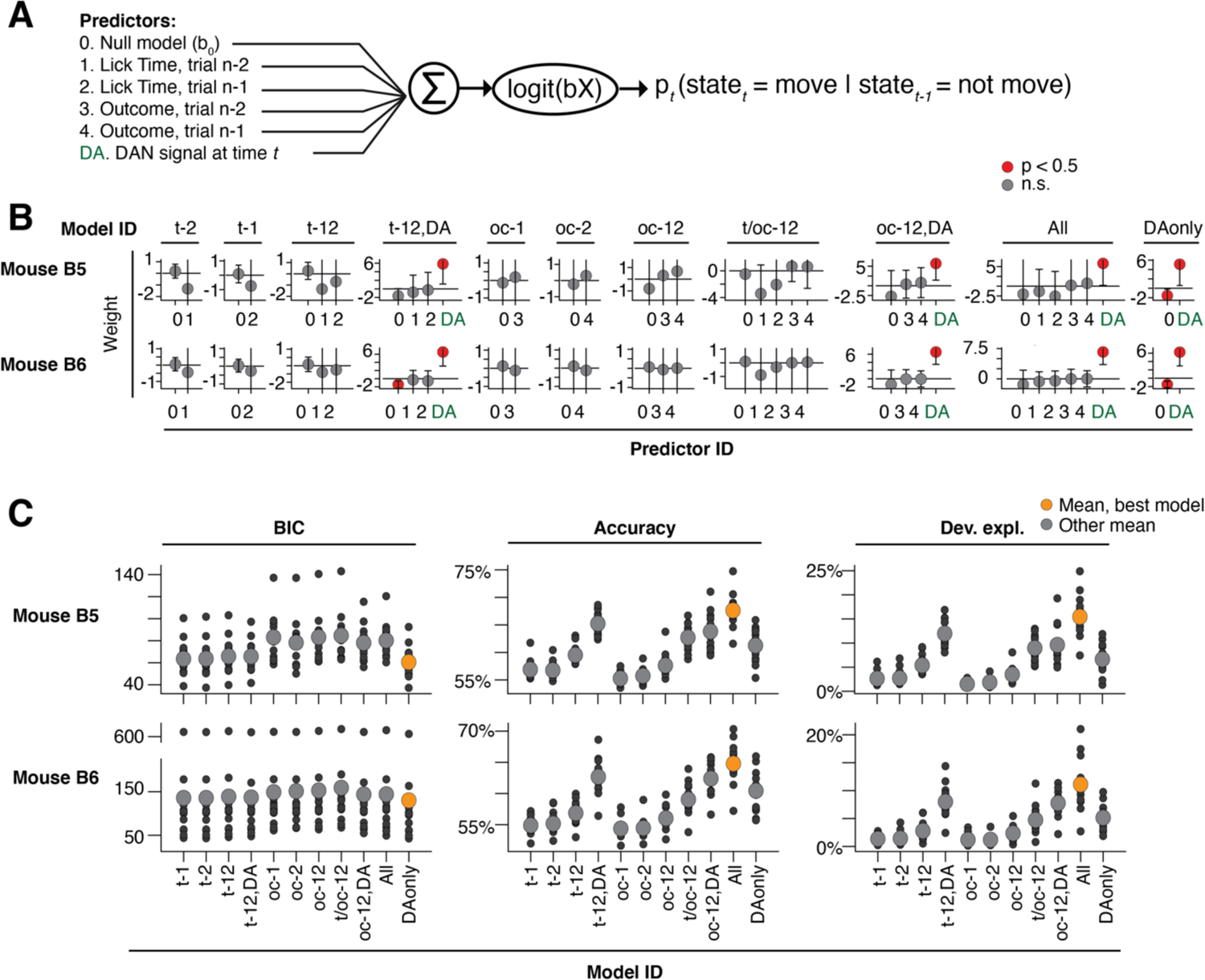
Probabilistic movement time decoding model: model selection. **(A)** Model schematic. To assess previous trial history on the same footing as dopaminergic signals, time *t* during model selection was limited to a 500 ms “time-slice,” with each time-slice fit separately by the model. Dopaminergic signals were averaged within each time-slice, such that each trial provided one and only one dopaminergic measurement, one set of trial history terms, and one movement state per time slice (see ***Methods: Single-trial probabilistic movement state decoding model, model selection***). **(B)** Model fit weights. Model ID: corresponds to the predictors included from the schematic. x-axis labels: the predictor ID from the schematic. Predictor weights averaged across time-slices. **(C)** Model selection criteria. The model omitting the previous trial history predictors (predictors #1-4) was consistently the best model as selected by BIC, AIC and AICc (results similar across metrics, BIC shown alone for clarity). Source data: Figure 8***—source data*.**

**Figure 8—figure supplement 2.**
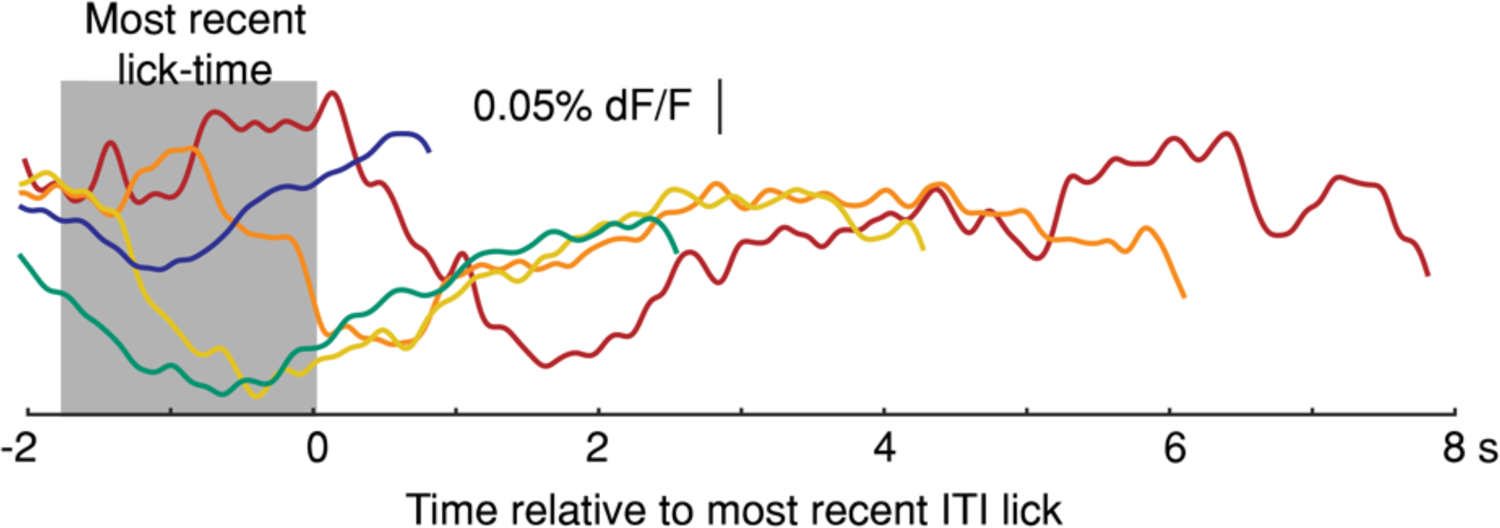
Average Intertrial Interval (ITI) GCaMP6f signals aligned to most recent previous lick-time. Signals plotted up to onset of next spontaneous, self-initiated lick during the ITI. (1 mouse, 5 sessions, truncated 150 ms before lick). Source data: Figure 8***—source data*.**

